# Mechanism of chaperone recruitment and retention on mitochondrial precursors

**DOI:** 10.1101/2025.01.18.633710

**Authors:** Szymon Juszkiewicz, Sew-Yeu Peak-Chew, Ramanujan S. Hegde

## Abstract

Nearly all mitochondrial proteins are imported into mitochondria from the cytosol. How nascent mitochondrial precursors acquire and sustain import-competence in the cytosol under normal and stress conditions is incompletely understood. Here, we show that under normal conditions, the Hsc70 and Hsp90 systems interact with and redundantly minimize precursor degradation. During acute import stress, Hsp90 buffers precursor degradation, preserving proteins in an import-competent state until stress resolution. Unexpectedly, buffering by Hsp90 relies critically on a mitochondrial targeting signal (MTS), the absence of which greatly decreases precursor-Hsp90 interaction. Site-specific photo-crosslinking and biochemical reconstitution showed how the MTS directly engages co-chaperones of Hsc70 (St13 and Stip1) and Hsp90 (p23 and Cdc37) to facilitate chaperone retention on the mature domain. Thus, the MTS has a previously unappreciated role in regulating chaperone dynamics on mitochondrial precursors to buffer their degradation and maintain import competence, functions that may facilitate restoration of mitochondrial homeostasis after acute import stress.

**Significance statement:**

- Mitochondrial proteins encoded by the nuclear genome are synthesized in the cytosol before their subsequent import into mitochondria. The factors that recognize mitochondrial precursors in the cytosol to maintain their import-competence are incompletely defined.
- Using a systematic site-specific photo-crosslinking strategy, the authors find that the mitochondrial targeting signal (MTS) is directly recognized by co-chaperones of Hsc70 and Hsp90. The co-chaperones facilitate recruitment, retention, and remodeling of these general chaperones on the nascent precursor protein.
- Chaperone retention becomes particularly important during mitochondrial stress, when precursors must avoid degradation during a prolonged period in the cytosol.

## Introduction

The majority of newly synthesized eukaryotic proteins are segregated from the cytosol to a membrane-enclosed compartment (Wickner and Schekman, 2005). Those destined for the ER lumen and mitochondrial matrix must be transported through translocation channels that can only accommodate unstructured linear polypeptides (Rapoport, 2007; Wiedemann and Pfanner, 2017). Thus, it is crucial that these nascent precursors avoid appreciable folding until they cross the membrane. However, unfolded proteins such as non-translocated precursors are typically targets for quality control pathways that initiate their degradation (Hessa *et al*., 2011; Rodrigo-Brenni *et al*., 2014; Itakura *et al*., 2016; Hegde and Zavodszky, 2019; Hipp *et al*., 2019). This means that ER and mitochondrial precursors must not only avoid folding, but also evade degradation until their transport.

One solution to this problem is to target the nascent protein to a translocation channel at an early stage in synthesis, allowing its co-translational transport across the membrane (Akopian *et al*., 2013). Whereas this is the major mechanism for ER translocation (Chartron *et al*., 2016; Costa *et al*., 2018; Guna and Hegde, 2018), most mitochondrial precursors are fully synthesized in the cytosol prior to their post-translational targeting and translocation (Wiedemann and Pfanner, 2017). This is supported by proximity-based ribosome profiling experiments (Williams *et al*., 2014), translocation assays in vitro (Young *et al*., 2003), pulse labelling assays in cells (Horwich *et al*., 1985), and the lack of appreciable membrane-bound ribosomes on the mitochondrial surface in electron microscopy studies (Palade, 1955). How mitochondrial precursors retain translocation competence and avoid degradation is incompletely understood (Wiedemann and Pfanner, 2017).

In the case of cytosol-localized ER precursors, the hydrophobic signal peptide is a degradation signal that avoids recognition by quality control because the targeting factor SRP can shield it shortly after it emerges from the ribosome (Hessa *et al*., 2011). This results in a hierarchy where targeting by SRP is the first option, the failure of which results in binding by other signal-binding factors such as Bag6 or the Ubiquilin family proteins (Hessa *et al*., 2011; Itakura *et al*., 2016). These quality control factors recruit ubiquitin ligases that target the bound substrate for degradation (Rodrigo-Brenni *et al*., 2014). A similar two-tiered hierarchy operates on tail-anchored (TA) membrane proteins, with the targeting signal (in this case, the hydrophobic C-terminal transmembrane domain) serving as a degron when recognition by the targeting factor fails (Hessa *et al*., 2011; Itakura *et al*., 2016; Shao *et al*., 2017). The role of the amphipathic mitochondrial targeting sequence (MTS), if any, in the key triage decision between translocation and degradation of mitochondrial matrix protein precursors is not known.

The main factors implicated in maintaining the translocation competence of mitochondrial matrix precursors are Hsc70 and Hsp90, general cytosolic chaperones (Neupert and Pfanner, 1993; Komiya *et al*., 1996; Young *et al*., 2003; Fan *et al*., 2006). These presumably bind dynamically along the polypeptide to impede folding, although the molecular details are not clear. In addition, crosslinking and interaction studies have suggested that the MTS is recognized by specific putative targeting factors (Murakami and Mori, 1990; Ono and Tuboi, 1990a, 1990b; Murakami *et al*., 1992; Hachiya *et al*., 1993, 1994, 1995), although the identity of these proteins was either never determined or could not be convincingly validated with recombinant proteins (Alam *et al*., 1994). If the MTS fails to initiate translocation, it could plausibly serve as a degron given its partial hydrophobic character, by analogy to ER-destined precursors and TA proteins. Indeed, recent work has implicated the MTS as a degron recognized by the SIFI complex, comprised of the ubiquitin ligases UBR4 and KCMF1, the E2 enzyme UBE2A, and calmodulin (Grabarczyk *et al*., 2024; Haakonsen *et al*., 2024; Yang *et al*., 2024).

The triage problem becomes particularly acute during mitochondrial stress, when import of precursors is markedly impaired (Wang and Chen, 2015; Wrobel *et al*., 2015; Weidberg and Amon, 2018). While these unimported precursors must eventually be degraded, there may be a benefit to delaying degradation in case the stress resolves quickly. This would allow precursor import to replenish the mitochondria of crucial proteins without having to re- synthesize them. Achieving the correct balance between precursor retention versus degradation is likely to be important for both cytosolic and mitochondrial homeostasis. Prolonged cytosolic retention risks aggregation and chaperone sequestration, whereas aggressive degradation risks eliminating import-competent proteins. How cells balance the retention of translocation competence with degradation, the role of the MTS in this triage reaction, and the mechanisms and factors involved are all largely unclear.

To begin addressing this issue, we investigated the suite of cytosolic interactors of the mammalian ornithine transcarbamylase precursor (pOTC), a model mitochondrial matrix enzyme. Although pOTC engages the Hsc70 and Hsp90 chaperones as expected, these do not interact with the MTS. Instead, the MTS interacts directly with Hsc70 and Hsp90 co- chaperones St13, p23, and Cdc37. These co-chaperones facilitate prolonged retention of their chaperones on the mature domain, thereby maintaining import competence. This function becomes particularly crucial during acute import stress, where the MTS impedes precursor degradation in a Hsp90-dependent manner. This function allows rapid import of retained precursors upon stress resolution, which may aid in restoration of mitochondrial homeostasis.

## Results

### Hsc70 and Hsp90 buffer pOTC degradation in the cytosol

To characterize the early steps of mitochondrial precursor biogenesis, we translated human pOTC in rabbit reticulocyte lysate, an import-competent in vitro translation system devoid of mitochondria (Horwich *et al*., 1985; Söllner *et al*., 1991). Affinity purification of newly synthesized pOTC under native conditions via its C-terminal twin-strep tag (TST) recovered Hsc70, Hsp90, and their co-chaperones as major, specific, and near-stoichiometric interaction partners (Fig. 1A, Table S1). This is consistent with the long-standing model that Hsc70 and Hsp90 retain mitochondrial precursors in an unfolded state to facilitate subsequent precursor import (Neupert and Pfanner, 1993).

**Figure 1.**
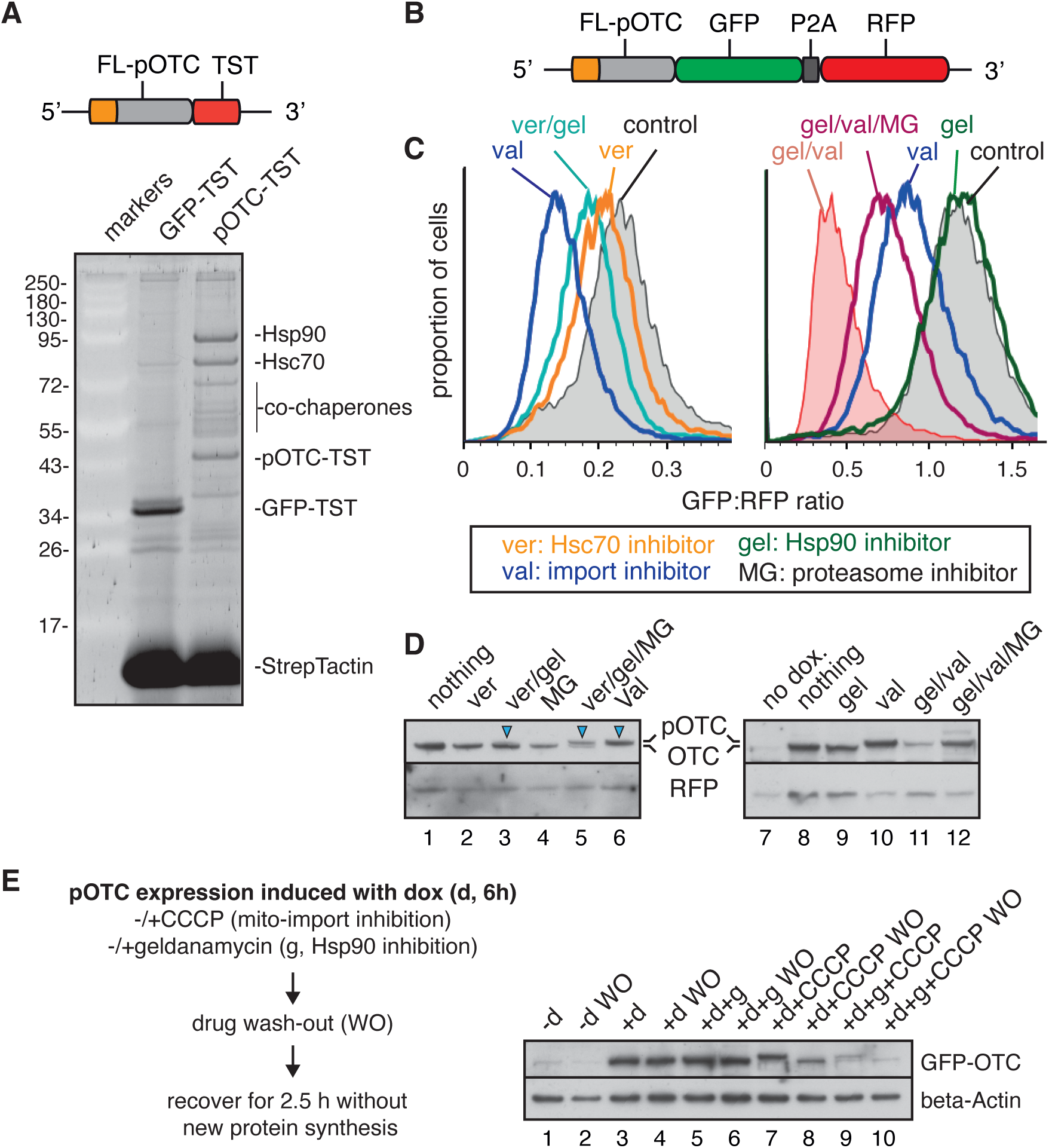
Hsp70 and Hsp90 buffer pOTC degradation in the cytosol. **(A)** Mitochondrial precursor of full length OTC (FL-pOTC) was translated in RRL and purified under native cond- tions via a C-terminal Twin-Strep tag (TST). The proteins eluted from the resin with biotin were analyzed by SDS-PAGE (shown below the diagram) and quantitative mass spectrometry (see Table 1). TST-tagged GFP served as a negative control. **(B)** Diagram of the reporter construct used to monitor stability and import of full length pre-OTC (FL-pOTC, with the MTS in orange). **(C, D)** Expression of the stably integrated reporter depict- ed in (B) was induced in HEK 293 Trex cells in the presence of 100 nM mitochondrial import inhibitor valinomy- cin (val) and 2 μM of Hsp90 inhibitor geldanamycin (gel) or 30 μM of Hsp70 inhibitor VER-155008 (ver), and/or 10 μM proteasome inhbitor MG132 for 7h. **(C)** Cells were then analyzed by flow cytometry or **(D)** immunoblot- ting. Histograms representing GFP fluorescence corrected for the expression of RFP and individual OTC-GFP and RFP blots are shown. Note the GFP:RFP ratio is relative and should only be used to compare conditions within a single experiment, but not between the two experiments. **(E)** Expression of the OTC-GFP reporter was induced in the presence or absence of 2 μM gel and/or 10 μM mitochondrial import inhibitor CCCP for 6h. Then, the drugs were washed out and cells were allowed to recover for 2.5h in the presence of 100 μg/ml protein synthesis inhibitor cycloheximide before lysis and analysis by immunoblotting.

**Table 1.** Constructs and antibodies used in this study.

| Name | Description and usage | Reference |
| --- | --- | --- |
| **Constructs** |  |  |
| ΔMTS-pOTC-TST | gBlock, sp64-based expression cassette for in vitro translation | This study |
| FL-pOTC-TST | gBlock, sp64-based expression cassette for in vitro translation | This study |
| FL-MRPS15-TST | gBlock, sp64-based expression cassette for in vitro translation | This study |
| ΔMTS-MRPS15-TST | gBlock, sp64-based expression cassette for in vitro translation | This study |
| GFP-TST | gBlock, sp64-based expression cassette for in vitro translation | This study |
| F3amber-FL-pOTC-TST | gBlock, sp64-based expression cassette for in vitro translation and site-specific photo-crosslinking | This study |
| R6amber-FL-pOTC-TST | gBlock, sp64-based expression cassette for in vitro translation and site-specific photo-crosslinking | This study |
| L9amber-FL-pOTC-TST | gBlock, sp64-based expression cassette for in vitro translation and site-specific photo-crosslinking | This study |
| A12amber-FL-pOTC-TST | gBlock, sp64-based expression cassette for in vitro translation and site-specific photo-crosslinking | This study |
| G17amber-FL-pOTC-TST | gBlock, sp64-based expression cassette for in vitro translation and site-specific photo-crosslinking | This study |
| H18amber-FL-pOTC-TST | gBlock, sp64-based expression cassette for in vitro translation and site-specific photo-crosslinking | This study |
| M21amber-FL-pOTC-TST | gBlock, sp64-based expression cassette for in vitro translation and site-specific photo-crosslinking | This study |
| N24amber-FL-pOTC-TST | gBlock, sp64-based expression cassette for in vitro translation and site-specific photo-crosslinking | This study |
| A12amber-FL-pOAT-TST | gBlock, sp64-based expression cassette for in vitro translation and site-specific photo-crosslinking | This study |
| L9amber-FL-MRPS15-TST | gBlock, sp64-based expression cassette for in vitro translation and site-specific photo-crosslinking | This study |
| V17amber-FL-MRPS15-TST | gBlock, sp64-based expression cassette for in vitro translation and site-specific photo-crosslinking | This study |
| I15amber-FL-pCOXIV1 | gBlock, sp64-based expression cassette for in vitro translation and site-specific photo-crosslinking | This study |
| px459-ST13-sgRNA1 | Plasmid, CRISPR-Cas9-mediated knockout of ST13 in mammalian cells | This study |
| pCMV-ST13-1xFLAG | Plasmid, transient overexpression of recombinant ST13 in mammalian cells | This study |
| pCMV-ΔTPR-ST13-1xFLAG | Plasmid, transient overexpression of recombinant ST13 in mammalian cells | This study |
| pCMV-ΔSTI-ST13-1xFLAG | Plasmid, transient overexpression of recombinant ST13 in mammalian cells | This study |
| pCMV-F335amber-ST13-1xFLAG | Plasmid, transient overexpression of recombinant ST13 using amber suppression in mammalian cells | This study |
| pCMV-Q336amber-ST13-1xFLAG | Plasmid, transient overexpression of recombinant ST13 using amber suppression in mammalian cells | This study |
| pCMV-M345amber-ST13-1xFLAG | Plasmid, transient overexpression of recombinant ST13 using amber suppression in mammalian cells | This study |
| pCMV-N356amber-ST13-1xFLAG | Plasmid, transient overexpression of recombinant ST13 using amber suppression in mammalian cells | This study |
| pcDNA5-FL-pOTC-GFP-P2A-mCherry | Plasmid, stable integration of the dual fluorescence reporter into the Flp-In Trex cells | This study |
| pAS-Pyl-AF | Plasmid, transient overexpression of the *Methanosarcina mazei* pyrrolysyl-tRNA synthetase carrying Y306A and Y384F mutations and tRNAPylCUA pair | (O'Donnell *et al.*, 2020) |
| **Antibodies** |  |  |
| Anti-St13 | #8014, clone 10B7-C3, mouse monoclonal (used for western blotting) | Cell Signaling Technology |
| Anti-St13 | #26581-1-AP, rabbit polyclonal (used for IP) | Proteintech |
| Anti-Stip1 | #15218-1-AP, rabbit polyclonal | Proteintech |
| Anti-p23 | #A304-499A, rabbit polyclonal | Bethyl |
| Anti-Cdc37 | #A302-489A, rabbit polyclonal antibody | Bethyl |
| Anti-Fkbp5 | #A301-430A, rabbit polyclonal | Bethyl |
| Anti-DnajA2 | #12236-1-AP, rabbit polyclonal | Proteintech |
| Anti-Btf3 | #A302-319A, rabbit polyclonal | Bethyl |
| Anti-Srp54 | #610941, mouse monoclonal | BD Biosciences |
| Anti-Hsc70/Hsp73 | #ADI-SPA-815, clone 1B5, rat monoclonal | Enzo Life Sciences |
| Anti-Hsp90alpha | #ab13495, rabbit polyclonal | Abcam |
| Anti-FLAG | #A8592, HRP-conjugated, mouse monoclonal | Sigma-Aldrich |
| Anti-StrepTagII | #ab76949, rabbit polyclonal | Abcam |
| Anti-beta-Actin | #A3854, HRP-conjugated, mouse monoclonal | Sigma-Aldrich |
| Anti-GFP | Homemade, rabbit polyclonal | (Chakrabarti and Hegde, 2009) |
| Anti-RFP | Homemade, rabbit polyclonal | (Chakrabarti and Hegde, 2009) |

To test the functional relevance of the Hsc70 and Hsp90 systems, we used a dual-fluorescence ratiometric assay (Itakura *et al*., 2016; Chitwood *et al*., 2018) for monitoring the stability and import of pOTC. In this assay, a reporter encoding pOTC tagged with GFP at the C-terminus is followed by a P2A ribosome skipping sequence and RFP (Fig. 1B). The reporter is stably integrated into a single doxycycline-inducible locus in HEK293 T-REx cells. The GFP:RFP ratio monitored by flow cytometry reports on the stability of pOTC-GFP relative to the RFP translation control. Induction of reporter expression (for 7 h) in the presence of the Hsc70 inhibitor VER-155008 (Ver) (Williamson *et al*., 2009) resulted in a modest reduction in pOTC-GFP relative to untreated cells (Fig. 1C). This reduction was further enhanced by combining Ver with the Hsp90 inhibitor geldanamycin (Gel) (Whitesell *et al*., 1994), whereas Gel alone had no obvious effect.

Parallel immunoblotting of these samples showed a small amount of pOTC (which migrates slightly slower than mature OTC) in cells treated with Ver/Gel (Fig. 1D, lane 3, arrowhead). Concomitant treatment with the proteasome inhibitor MG132 (MG) (Lee and Goldberg, 1998) increased pOTC levels in Ver/Gel-treated cells (lane 5), but not untreated cells (lane 4). The amount of precursor in Ver/Gel/MG-treated cells was roughly twice that of mature OTC, indicating that simultaneous inhibition of Hsc70 and Hsp90 impairs most but not all pOTC import. By contrast, dissipating the mitochondrial membrane potential with valinomycin (Val) (Martin *et al*., 1991) completely inhibited pOTC import (as seen by immunoblotting; Fig. 1D, lane 6) and led to substantial pOTC degradation (as monitored by the GFP:RFP ratio; Fig. 1C).

These results are consistent with a model where the Hsc70 and Hsp90 systems directly engage pOTC and redundantly facilitate its mitochondrial import. The lack of effect of Gel alone on pOTC levels suggests that under normal conditions, the Hsp90 system has little or no role in import, presumably because the Hsc70 system serves this role. Surprisingly, inhibiting Hsp90 during mitochondrial import stress sharply reduced pOTC levels relative to import stress alone (Val compared to Gel/Val; Fig. 1C). This could be ascribed to degradation because pOTC could be partially stabilized by simultaneously inhibiting the proteasome (Val/Gel/MG). This conclusion was verified by parallel immunoblotting (Fig. 1D, lanes 10-12). Here, one can appreciate that during import stress, pOTC accumulates in the cytosol with no obvious degradation (presumably because immunoblotting is not especially quantitative, unlike the GFP:RFP ratio measured by flow cytometry). By contrast, pOTC is degraded efficiently if Hsp90 is simultaneously inhibited.

Similar results were seen with CCCP (Fig. 1E), another agent that dissipates the mitochondrial membrane potential (Martin *et al*., 1991). Furthermore, cytosolic pOTC that accumulates under import stress (Fig. 1E, lane 7) is competent for import and is converted to OTC when CCCP is washed out (WO; Fig. 1E, lane 8). In the presence of Hsp90 inhibition, however, very little pOTC remains during the CCCP treatment due to degradation (Fig. 1E, lane 9). The residual amount of pOTC seen with CCCP/Gel does not seem to be import competent as no appreciable increase in mature OTC was seen upon washout of the inhibitors (Fig. 1E, lane 10). These results indicate that cytosolic precursors are maintained in an import-competent state by a combination of the Hsc70 and Hsp90 systems. Under normal conditions of active import, the brief residence time in the cytosol seems to be managed primarily by Hsc70, whereas prolonged cytosolic residence during import stress relies on Hsp90.

### Hsp90 broadly buffers mitochondrial precursor degradation during import stress

To determine whether Hsp90-mediated stabilization of mitochondrial precursors during mitochondrial import stress is a general phenomenon, we performed Tandem-Mass-Tag (TMT) quantitative proteomics. We analyzed cytosol from cells in which we inhibited mitochondrial import with or without simultaneous inhibition of Hsp90 and compared them to the cytosol of untreated cells (Fig. 2A). All of these cells were expressing the pOTC-GFP reporter, whose behavior under these conditions was known from the preceding experiments and served as a positive control. We detected ∼5500 proteins, of which ∼550 were annotated as mitochondrial proteins (marked as blue and red dots) (Fig. 2B, C; Fig. S1; Table S2).

**Figure 2.**
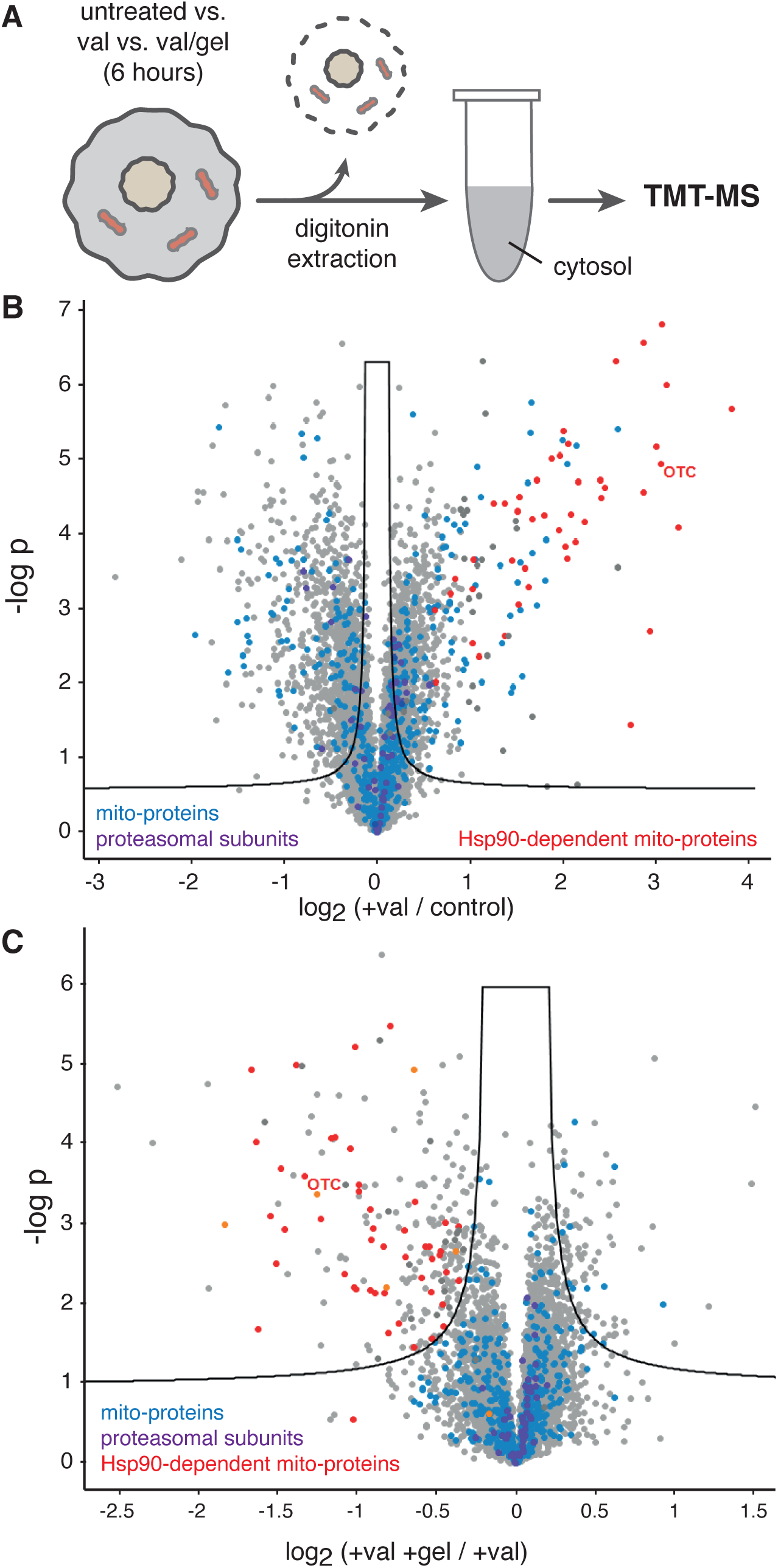
Hsp90 broadly buffers mitochondrial precursor degradation during import stress. (A) Schematic representation of the experimental setup. Digitonin-extracted cytosolic extracts from cells expressing OTC-GFP reporter in the presence or absence of mitochondrial import (val) or Hsp90 activity (gel) for 6 h were analyzed by quantitative mass spectrometry using TMT. The identified proteins were plotted by log_2_ fold difference (x axis) against -log p value. **(B)** Comparison of protein abundance in cytosolic extracts from control cells and cells experiencing mitochondrial import stress (val). **(C)** Comparison of protein abundance in cells under mitochondrial import stress with or without an active Hsp90 system. Mitochondrial proteins are depicted in blue, proteasomal subunits in purple and mitochondrial-proteins destabilized after Hsp90 inhibition are in red. The black lines on both plots set 0.05 FDR. See also Fig. S1 and Table S2.

As expected, the amount of cytosolic OTC was increased with import stress (Fig. 2B), an effect that was largely reversed by simultaneous Hsp90 inhibition (Fig. 2C). We found ∼50 additional proteins that showed the same Hsp90-dependent behavior. Almost all of these proteins were mitochondrial matrix proteins or inner mitochondrial membrane proteins (marked as red dots). Importantly, mitochondrial import inhibition in our experiment did not induce any significant changes to the cytosolic proteome. In particular, no changes in proteasome abundance were detected (individual subunits marked with purple dots). This contrasts with yeast cells that upregulate the proteasome with chronic mitochondrial import stress (Wang and Chen, 2015; Wrobel *et al*., 2015).

Together with the focused analysis of pOTC (Fig. 1), the proteomic analysis in Fig. 2 shows that many mitochondrial precursors rely on Hsp90-mediated stabilization during mitochondrial import stress. In the case of pOTC, the stabilized cytosolic precursor retains import competence (Fig. 1E), suggesting that the other precursors are likely to be similar. Prolonged retention of precursors in an import competent state during mitochondrial stress would allow mitochondria to begin replenishing their proteome rapidly after stress resolution rather than relying on the slower process of de novo protein production. Because this key function relies on Hsp90, we turned our attention to the mechanism of Hsp90-mediated precursor stabilization.

### The MTS facilitates Hsp90 retention on mitochondrial precursors

Almost all of the Hsp90-dependent proteins detected in our proteomics experiment (tab 2 in Table S2) contain a predicted cleavable MTS, just like our model protein pOTC. We therefore investigated whether the MTS might play a role in Hsp90 retention. To dissect the molecular details of precursor-Hsp90 interactions, we turned to the in vitro translation system. As shown previously (Conboy and Rosenberg, 1981; Horwich *et al*., 1985), pOTC-TST produced in reticulocyte lysate was imported into mitochondria as judged by MTS cleavage (Fig. 3A). No impairment of pOTC import was seen in the presence of Hsp90 inhibition (Fig. 3A, lane 5), consistent with the data in cells (Fig. 1C-E), presumably due to other chaperones such as Hsc70 in the system.

**Figure 3.**
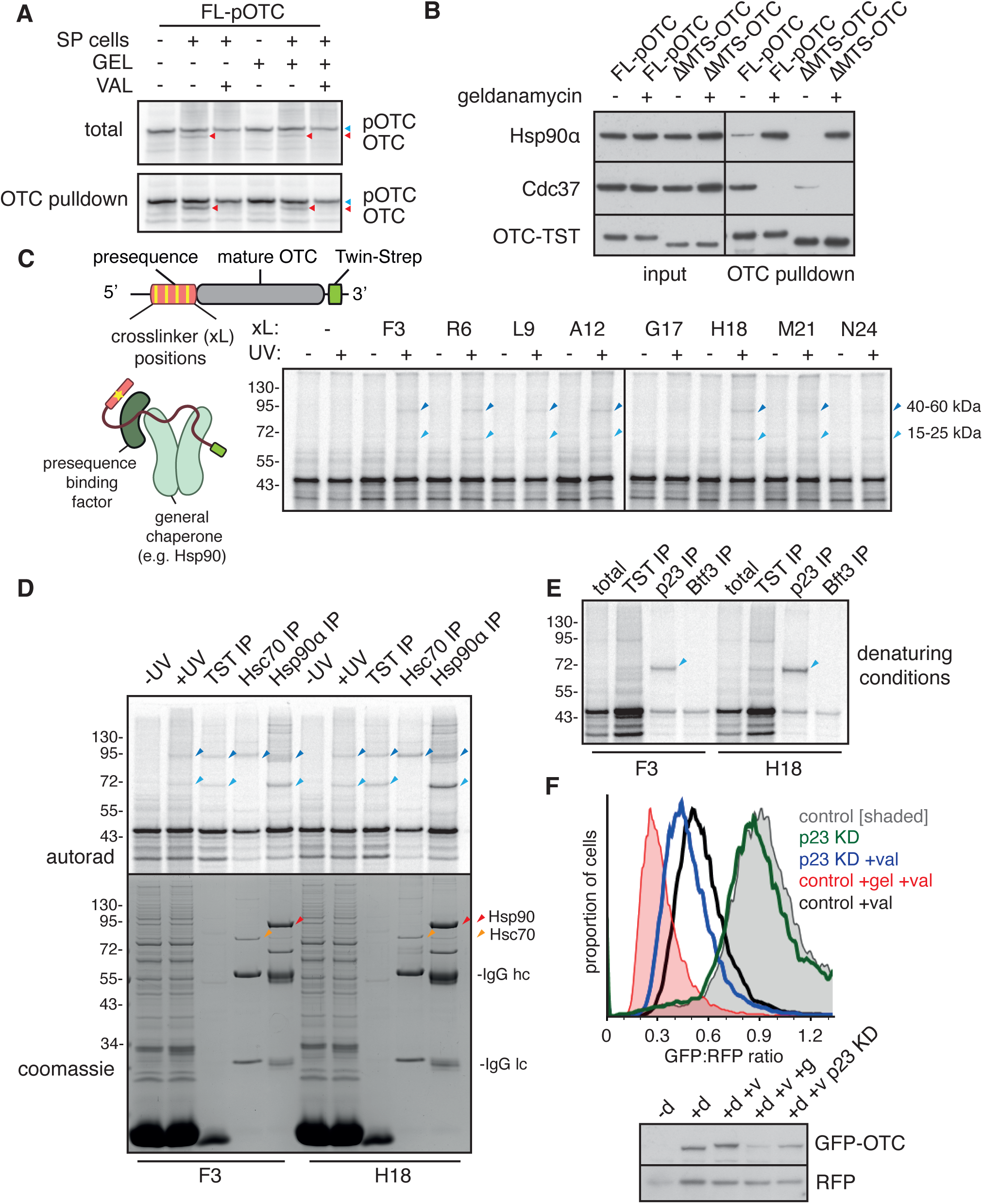
The MTS facilitates Hsp90 retention on mitochondrial precursors. (A) pOTC-TST was translated in RRL in the presence or absence of semi-permeabilized (SP) cells and valino- mycin (1 μM) and/or geldanamycin (2 μM). Total (top) or affinity purified (bottom) samples were analysed by SDS-PAGE and autoradiography. pOTC is marked with blue arrows, whereas mature OTC with cleaved MTS is marked with red arrows. Note that proteasome-mediated degradation is not active in the RRL system, which is why geldanamycin treatment does not lead to destabilization of the precursor as seen in cells. **(B)** Full length (FL) or ΔMTS OTC was translated in RRL in the presence or absence of 2 μM geldanamycin and affini- ty purified under native conditions via TST. Eluted samples were analyzed by immunoblotting alongside input samples. **(C)** Strategy to identify MTS-binding factors. Photo-activatable crosslinking amino-acid BpA was introduced into different positions within the MTS of pOTC using amber suppression. Amber mutants were translated in RRL and crosslinking was induced with UV. Crosslinking products were analyzed by SDS-PAGE. Two major crosslinking bands representing a 15-25 kDa adduct (light blue arrows) and a 40-60 kDa adduct (dark blue arrows) are marked. **(D)** pOTC with BpA at positions F3 or H18 was produced in RRL and crosslinked with UV. Samples were then immunoprecipitated with antibodies against Hsc70 or Hsp90α under native conditions and analyzed by SDS-PAGE. Major crosslinking bands are marked with the same colours as in (C). Coomassie stained gel (bottom) verifies efficient pulldown of Hsc70 (orange arrows) and Hsp90 (red arrows). **(E)** BpA-containing pOTC mutants were produced in RRL and crosslinked. Fully denatured samples were then immunoprecipitated with antibodies against p23 and an unrelated protein of similar size (Btf3), and analyzed by SDS-PAGE. **(F)** Cells with stably integrated pOTC-GFP-P2A-RFP reporter were treated with control siRNA or siRNA targeting p23 for 3 days and expression of the reporter was induced for another 7h in the presence of indicated drugs. Cells were then analyzed by flow cytometry (top) or immunoblotting (bottom). Histogram representing GFP:RFP ratio is shown. See also Figure S2.

In vitro translated pOTC-TST interacted with Hsp90, and its co-chaperone Cdc37, by co-IP, whereas these interactions were markedly reduced for ΔMTS-OTC-TST (Fig. 3B). For both pOTC-TST and ΔMTS-OTC-TST, the Hsp90 interaction was stabilized by geldanamycin, which also led to a complete loss of Cdc37 interaction. A similar MTS-dependent Hsp90 interaction was seen for MRPS15, an unrelated mitochondrial matrix precursor (Fig. S2A).

To understand this MTS-dependent Hsp90 interaction with mitochondrial precursors, we turned to site-specific photo-crosslinking. In this experiment, we probed the molecular environment of the MTS using p-Benzoyl-L-phenylalanine (BpA), a UV-activated crosslinker incorporated into each of eight different positions along the MTS via amber suppression (Lin *et al*., 2020). Surprisingly, none of the prominent crosslinking partners from any of the positions matched the size of Hsp90 (or Hsc70). Instead, the two primary crosslinking adducts corresponded to proteins of ∼15-25 kDa and ∼40-60 kDa (Fig. 3C). Given the similar crosslinking pattern from all positions (albeit at somewhat different ratios of the adducts), subsequent experiments focused on positions 3 and 18.

Although neither Hsc70 nor Hsp90 were direct MTS interactors, native IPs using antibodies against either chaperone could recover different subsets of the primary crosslinking products (Fig. 3D). The lower molecular weight adduct was selectively co-precipitated with Hsp90, whereas the higher molecular weight adduct co-precipitated with both Hsc70 and Hsp90. Depletion of ATP with apyrase prior to UV irradiation resulted in loss of the small adduct, but not the larger adduct (Fig. S2B). A major ATP-dependent interactor of Hsp90 in this size range is p23 (also known as PTGES3) (McLaughlin *et al*., 2006; Noddings *et al*., 2022), which acts to slow ATP hydrolysis by Hsp90. The identity of the lower crosslink as p23 was confirmed by denaturing IP (Fig. 3E). Crosslinking between p23 and the MTS was also seen for pOAT and MRPS15, two unrelated mitochondrial precursors (Fig. S2C, D).

To test the importance of p23 interaction with mitochondrial protein precursors, we acutely knocked down p23 in HEK cells using siRNA and analyzed our pOTC-GFP-P2A-RFP reporter. Depletion of p23 under steady-state conditions did not substantially impact the stability or import of pOTC (Fig. 3F; Fig. S2E). However, when we inhibited mitochondrial import with valinomycin in cells lacking p23, we observed a decrease in cytoplasmic pOTC-GFP. This phenotype partially mimics Hsp90 inhibition, indicating that p23 is important for the efficient buffering of mitochondrial precursors by Hsp90 during mitochondrial import stress, presumably by slowing down the Hsp90 cycle (McLaughlin *et al*., 2006; Noddings *et al*., 2022) and effectively preventing the precursors from reaching their native folded state.

### Cdc37 and St13, co-chaperones of Hsp90 and Hsc70, interact with the MTS

To identify the higher molecular weight (∼40-60 kD) crosslinking adduct(s), we inspected our pOTC interactome (Fig. 1A; Table S1). The Hsp90 co-chaperones Fkbp5 and Cdc37 were among the most prominent interactors in this size range. The higher molecular weight crosslink could be immunoprecipitated under denaturing conditions with antibody against Cdc37 (Fig. 4A, top panel) but not Fkbp5 (Fig. 4B, top panel). Nonetheless, the crosslink co-purified with both Cdc37 and Fkbp5 under native conditions (Fig. 4A, 4B bottom panels). In both experiments, the larger crosslink was immunoprecipitated only when the crosslinker was at position F3, but not at position H18, suggesting that different proteins of similar size crosslink to the MTS from the two positions, with only the adduct from F3 being Cdc37.

**Figure 4.**
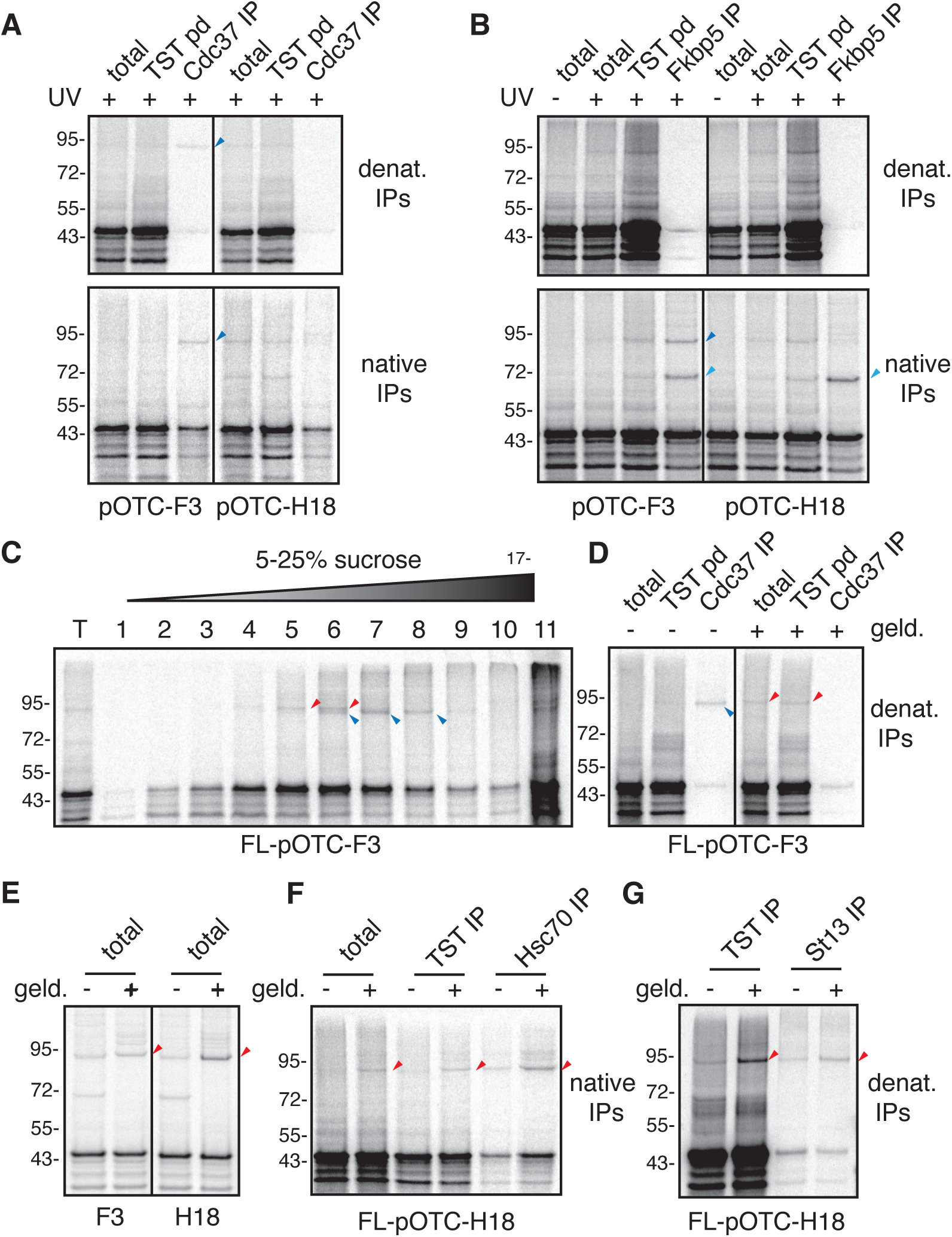
Cdc37 and St13, co-chaperones of Hsp90 and Hsc70, interact with the MTS. BpA-containing pOTC mutants were produced in RRL and crosslinked with UV. Fully denatured samples (top gels) or native samples (bottom gels) were then immunoprecipitated with antibodies against Cdc37 **(A)** or Fkbp5 **(B)** and analyzed by SDS-PAGE. **(C)** In vitro translated, BpA-containing pOTC was fractionated on a 5-25% sucrose gradient followed by UV crosslinking of individual fractions. Samples were analyzed by SDS-PAGE. Two independent crosslinks migrating close to each other on the gel are marked with red and blue arrows. **(D)** FL-pOTC containing BpA at position F3 was translated in vitro in the presence or absence of geldanamycin. Samples were then crosslinked with UV, immunoprecipitated with antibody against Cdc37 and analyzed by autoradiography. **(E)** FL-pOTC containing BpA at positions F3 or H18 were translated in vitro, crosslinked with UV and analyzed by autoradiography. **(F)** FL-pOTC-H18 was translated in vitro, and purified with antibody against Hsc70 under native conditions. **(G)** FL-pOTC-H18 was translated in vitro, crosslinked and purified with antibody against St13 under denaturing conditions.

Two closely-migrating adducts from position F3 could be separated by size fractionation through a sucrose gradient. One adduct (red arrows) migrated in fractions 4-6, whereas the other slightly smaller adduct (blue arrows) migrated deeper into the gradient in fractions 6-8 (Fig. 4C). When Hsp90 was inhibited with geldanamycin, the interaction (Fig. 3B) and crosslink to Cdc37, which proved to be the slightly smaller adduct, was lost (Fig. 4D), similarly to the p23 adduct (Fig 4E). Under these conditions, the other crosslink (red arrow), which was also seen from position 18, was retained (Fig. 4D, 4E). Native IP with anti-Hsc70 co-precipitated the geldanamycin-insensitive crosslink (Fig. 4F) suggesting that it might be St13 (also known as Hip) (Höhfeld *et al*., 1995). St13 is not only in the correct size range, but was also the most prominent Hsc70 interactor co-purified with pOTC (Table S1). Denaturing IP with antibody against St13 confirmed the crosslink’s identity and verified that its interaction with the MTS is enhanced in the presence of geldanamycin (Fig. 4G). St13 also crosslinked to the MTS of the unrelated mitochondrial precursor pCOXIVl1 (Fig. S3).

Together with the above results, we conclude that the MTS directly interacts with three co-chaperones (p23, Cdc37, and St13) and indirectly interacts with at least one (Fkbp5). Three of these are part of the Hsp90 system, with Cdc37 generally facilitating client loading onto Hsp90 (Stepanova *et al*., 1996), Fkbp5 facilitating client maturation (Pirkl and Buchner, 2001; Noddings *et al*., 2023), and p23 facilitating client retention on Hsp90 by slowing its ATPase activity (Schopf *et al*., 2017). The last co-chaperone, St13, similarly slows the Hsc70 ATPase cycle to promote its holdase activity (Höhfeld *et al*., 1995; Li *et al*., 2013). Notably, the St13- Hsc70 complex seems to be distinct from the Hsp90-containing complex given that they can be separated by size on a sucrose gradient (Fig. 4C). As expected, the latter has a larger native size given that Hsp90 forms a dimer. Furthermore, the enhanced interaction with St13 when Hsp90 is inhibited (Fig. 4E-G) raised the possibility that the two complexes may have a precursor-product relationship, with the Hsc70 complex engaging first before a handover to the Hsp90 system.

### St13 engages the MTS co-translationally and is retained after ribosome release

The Hsc70 system can interact with its clients co-translationally early during biosynthesis. To test whether St13 might engage the MTS as it emerges from the ribosome, we generated ribosome-nascent-chain complexes (RNCs) stalled ∼70 amino acids downstream of the MTS of pOTC (Fig. 5A). A photo-crosslinker at position H18 of the MTS in these RNCs detected several specific crosslinks (Fig. 5B, arrowheads). The crosslinks were unchanged with ATP depletion by apyrase, but only one crosslink (red arrowhead) remained and was enhanced upon dissociation of the ribosome with EDTA and RNase A (Fig. 5B). This suggests that an exposed MTC on an RNC is sampled by multiple factors, one of which becomes the dominant interaction partner upon release into the bulk cytosol.

**Figure 5.**
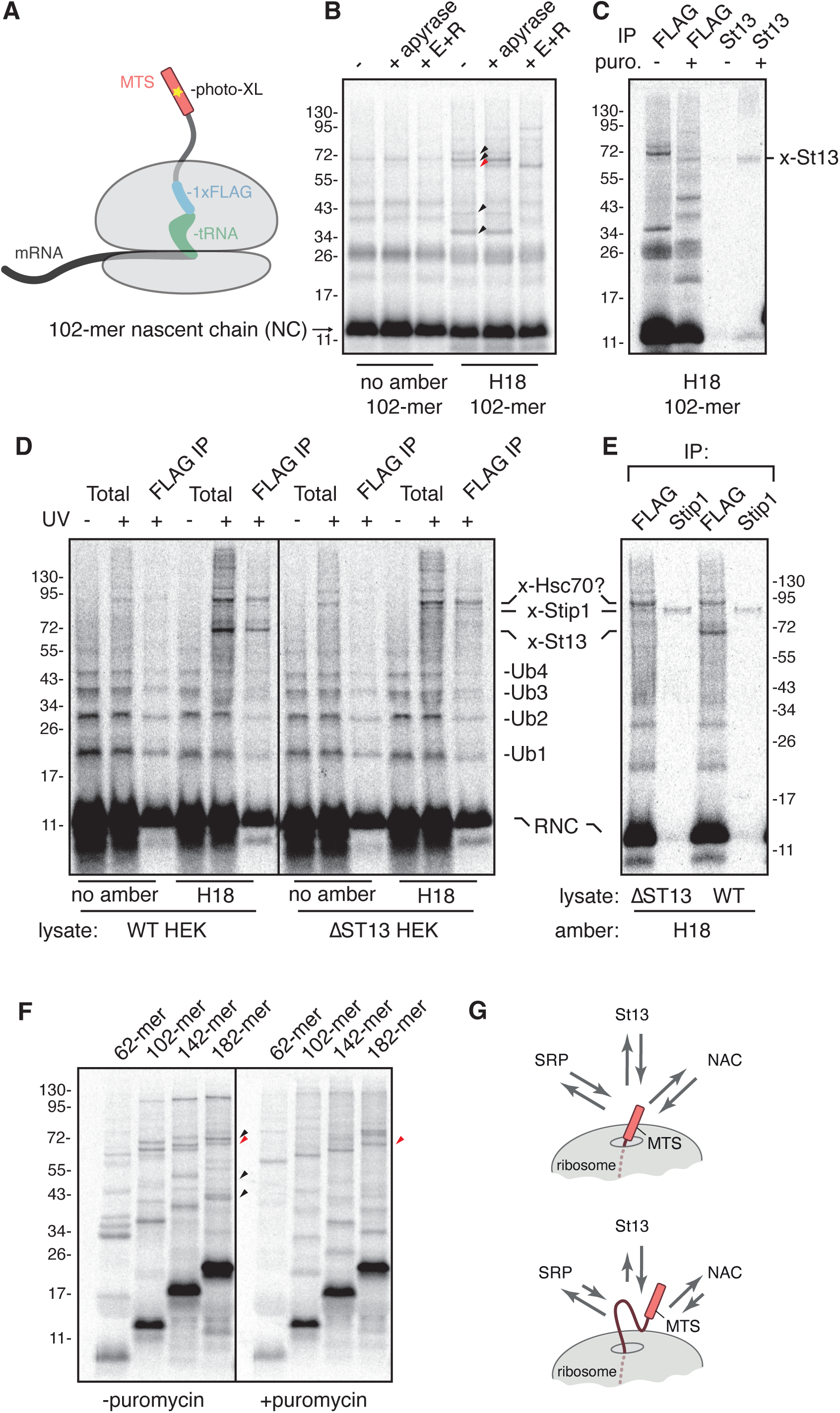
St13 engages the MTS co-translationally and is retained after ribosome release. (A) Experimental strategy used to identify MTS-interacting factors during early biogenesis of mitochondrial precursors. **(B)** Ribosome-nascent chain complexes (RNCs) with or without BpA at position H18 were produced in RRL. Samples were then treated with nothing, apyrase to deplete ATP, or EDTA and RNAse A (+E+R) to release the polypeptide from the ribosome before crosslinking with UV. All samples in this and subsequent experi- ments were analyzed on SDS-PAGE after RNase treatment to remove attached tRNA. Specific crosslinks are marked with arrows, with the red arrow indicating the product that is retained even after nascent chain release from the ribosome. **(C)** 102-mer RNCs with BpA at position H18 were produced in RRL, treated with puromycin where indicated to release the nascent chain from the ribosome, irradiated with UV, and subjected to immunopre- cipitation (IP) via the nascent chain (FLAG tag) or St13. **(D)** RNCs with or without BpA at position H18 were produced in RRL, isolated via centrifugation, resuspended in cytosolic lysate from HEK293 cells, incubated with puromycin to release nascent chains from ribosomes, and irradiated with UV. After UV crosslinking, all samples were denatured and either analyzed directly (total) or IP via the FLAG tag on the nascent chain. **(E)** As in panel D, but also including IP via Stip1. **(F)** RNCs of different length with BpA at position H18 were produced in RRL, incubated without or with puromycin to release nascent chains from ribosomes, then irradiated with UV. Crosslinks are indicated with arrowheads, with red indicating the crosslink to St13. **(G)** Schematic depicting early events during mitochondrial precursor biogenesis. An MTS emerging from the ribosome initially interacts with NAC and SRP, most likely due to proximity of these abundant ribosome-associating factors. However, as more polypeptide becomes exposed and the MTS is more flexibly tethered to the ribosome, the St13 interaction is progressively favoured to facilitate Hsc70 recruitment.

Based on sizes and ribosome-dependence, we speculated, then verified by IPs, that two of the crosslinks are the beta subunit of Nascent Polypeptide-Associated Complex (NAC) and the 54 kD subunit of Signal Recognition Particle (SRP) (Fig. S4A, S4B). Note that unlike bona fide ER-destined substrates (Powers and Walter, 1996), the MTS interaction with SRP is salt-sensitive, indicating that it is not stably engaged (Fig. S4C). One of the weak crosslinks could be recovered by IP using anti-DNAJA2, a co-chaperone of Hsc70 (Fig. S4B); by contrast, crosslinking to the Hsp90 co-chaperone p23 was not detectable in the total crosslinking sample and was barely detectable after enrichment by IP (Fig. S4A). The smallest of the crosslinks (∼20 kDa) is likely to be a core ribosomal protein as it was retained in RNCs isolated under high-salt conditions that strip non-ribosomal proteins (Fig. S4C). As expected, this crosslink is lost when the nascent chain is released from the ribosome (Fig. 5A, S4C).

The crosslink that is enhanced upon release from the ribosome was identified by IP to be St13 (Fig. 5C). When isolated RNCs were dissociated with puromycin into cytosolic lysates derived from HEK293 cells, a major crosslink matching the size of St13 was again observed (Fig. 5D), indicating that this interaction is not specific to reticulocyte lysate. The identity of this crosslink in HEK293 cell lysate was verified to be St13 by its loss in lysates from St13 knockout cells (ΔSt13). In addition to the St13 crosslink, two additional higher molecular weight crosslinked bands were seen. The larger and more prominent crosslink was only partially dependent on the presence of BPA (with crosslinking presumably occurring via favorably positioned aromatic residues) and represents crosslinking to Hsc70 based on its size, abundance, interaction with Hsc70 and co-chaperones by co-IP (Fig. 1A) and a separate denaturing IP experiment using anti-Hsc70 antibody (Fig. S4D). The lower molecular weight crosslink was shown by IP to be the St13 homologue Stip1 (Fig. 5E).

Crosslinking analysis of different lengths of RNCs showed that the crosslink to St13, which is hardly detectable when the MTS first emerges from the ribosome, becomes progressively more prominent for longer RNCs (Fig. 5F, red arrowhead). Release of these different length nascent chains from the ribosome showed that St13 interacts similarly with all of them. These results indicate that an MTS displayed on an RNC can sample various nascent chain interaction partners including NAC, SRP, and St13. At early stages of mitochondiral precursor translation, ribosome-associated factors such as SRP and NAC have priority presumably due to their proximity to the emerging MTS. However, as the nascent chain is elongated and more of the polypeptide becomes exposed, St13 can better engage the MTS (Fig. 5G). After release from the ribosome, the MTS primarily engages St13 in both reticulocyte lysate and HEK cell lysate. St13 interacts with Stip1 and Hsc70 as part of a larger chaperone complex (Frydman and Höhfeld, 1997). In the absence of St13, it seems that its homologue Stip1 is still able to engage the MTS. Because Stip1 can also engage Hsp90, these results suggest a route by which the Hsc70 and Hsp90 systems are recruited to the nascent chain via co-chaperone engagement of the MTS, probably beginning initially with St13 sampling of RNCs.

### Mechanism of MTS engagement by St13

St13 has two major structural domains – the TPR motif through which it interacts with Hsc70 and the STI1 domain, whose function is unknown but whose predicted structure houses a semi-hydrophobic groove that could bind amphipathic substrates such as an MTS (Li *et al*., 2013; Fry *et al*., 2021). Consistent with this idea, reconstitution of ΔST13 HEK293 cells with St13 lacking the STI1 domain (ΔSTI) diminished adducts with a photo-crosslinker-containing MTS, which was otherwise strong with recombinant wild type St13 (Fig. 6A, 6B). To determine the site on St13 that contacts the MTS, we introduced photo-crosslinkers by amber suppression into the STI1 domain of recombinant St13 and repeated the crosslinking experiment. Three residues that line the hydrophobic groove of the predicted STI1 domain (Fig. 6C) each crosslinked to the MTS-containing substrate, but a residue outside this groove (N356) did not (Fig. 6D). Thus, the hydrophobic groove of the STI1 domain of St13 directly engages the MTS of a mitochondrial precursor.

**Figure 6.**
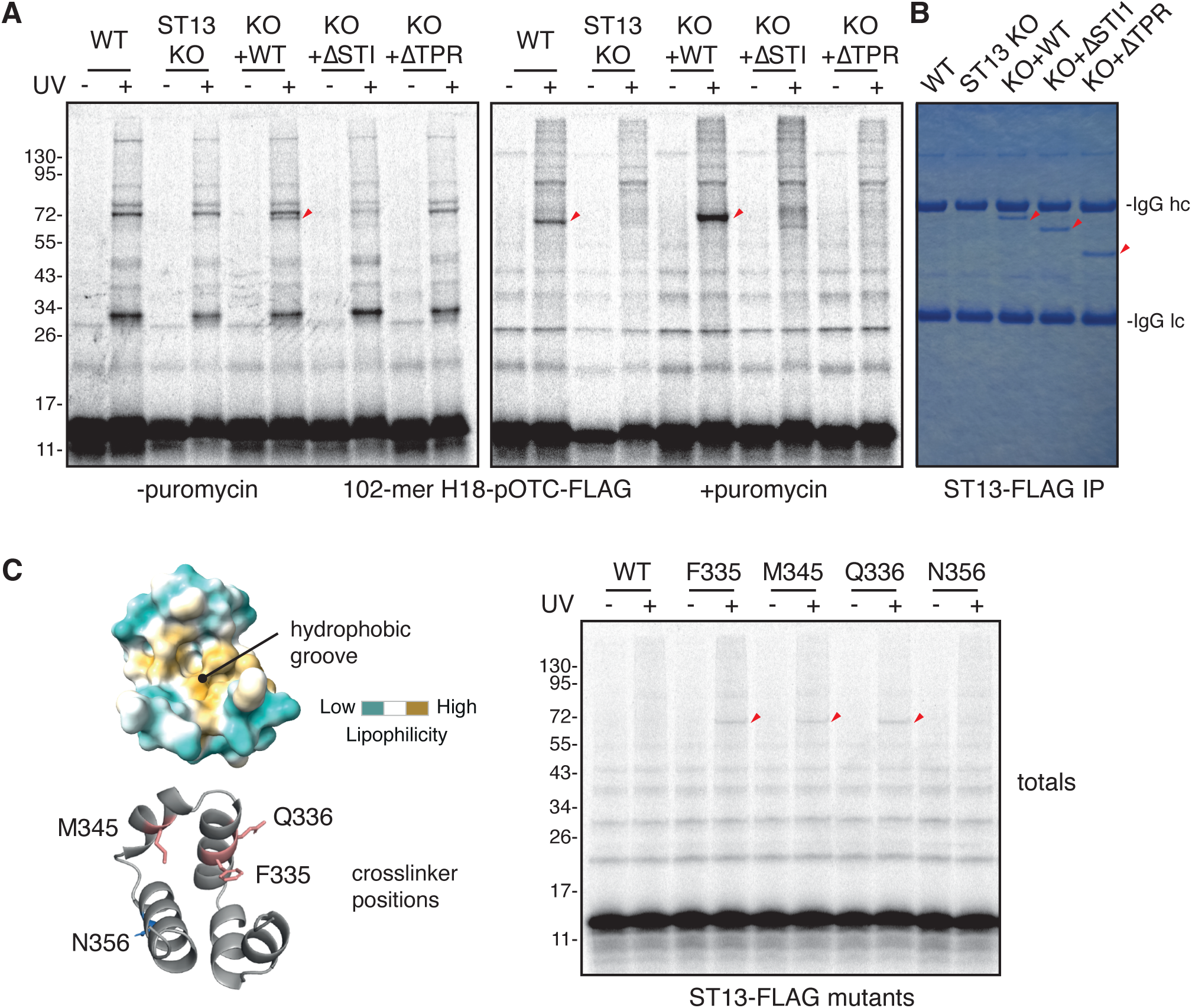
St13 engages MTS via its STI1 domain. (A) 102-mer ribosome nascent-chain complexes (RNCs) of pOTC with BpA at position H18 were synthesized in RRL, purified and resuspended in HEK lysates from WT or ΔST13 cells expressing the indicated St13 mutants, and crosslinked with UV. One set of nascent chains was UV-irradiated directly after addition of the lysates (left gel) whereas another set of nascent chains was released from ribosomes with puromycin before UV irradiation. Crosslinks between the radiolabelled pOTC and St13 are marked with red arrows. **(B)** Cytosolic extracts from (A) were denatured and immunoprecipitated with anti-FLAG antibodies to recover St13 mutants (marked with red arrows). Note that all mutants are expressed at the same level. **(C)** On the left, the STI1 domain of St13 as predicted by AlphaFold2, shown colored by hydrophobicity (top) and as a cartoon diagram (bottom). Positions at which the photo-activatable crosslinker AbK was introduced are marked. On the right, radiolabelled 102-mer RNCs of pOTC without any crosslinker were synthesized in RRL, purified via centrifugation, then resuspended in cytosolic extracts from ΔST13 cells transiently expressing ST13 mutants without AbK (WT) or with AbK at indicat- ed positions. Crosslinking was induced with UV. Crosslinks between the pOTC NC and ST13 mutants are marked with red arrows.

**Figure 7.**
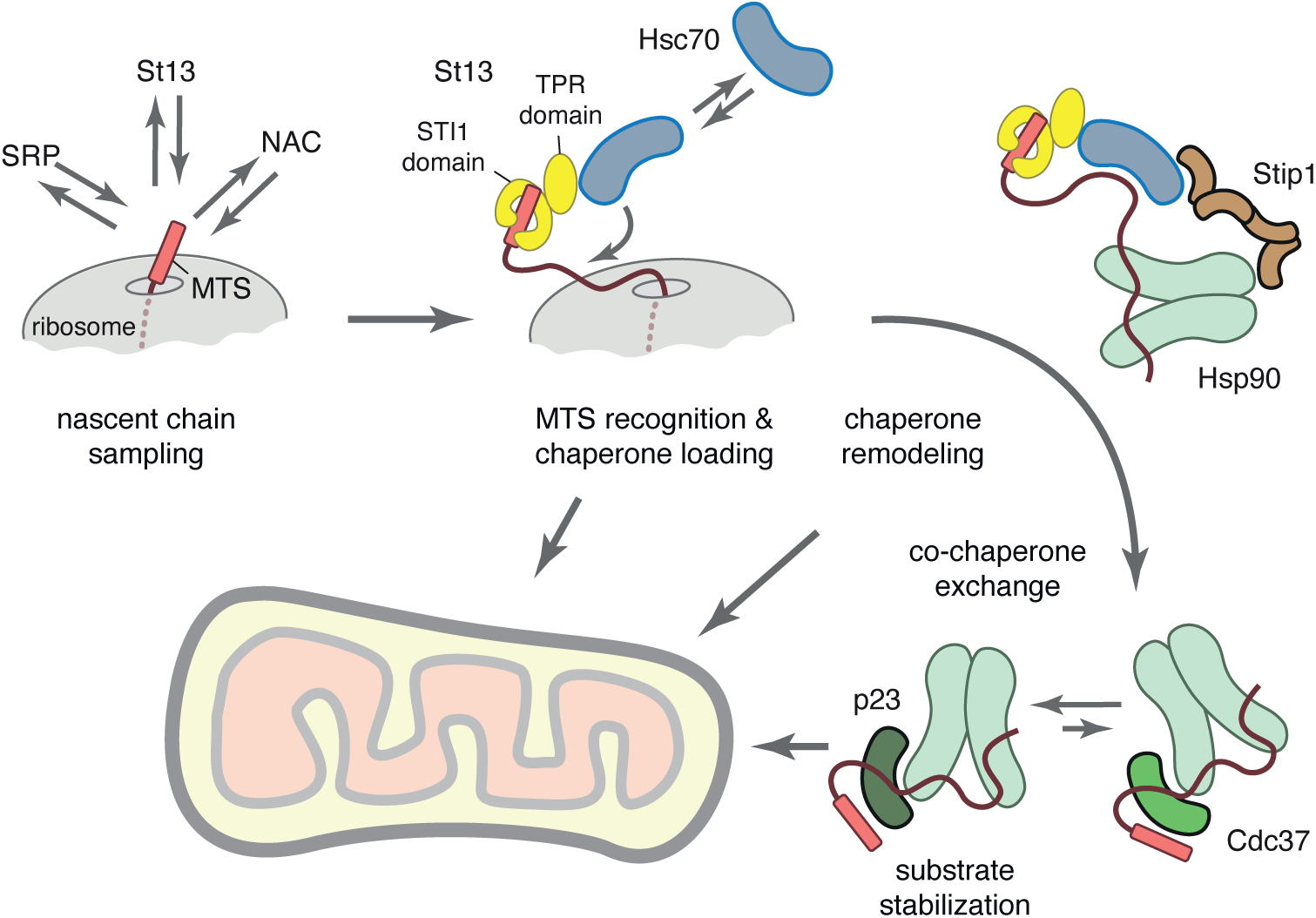
Cytosolic events during mitochondrial precursors’ biogenesis. MTS-containing mitochondrial precursors are sampled by multiple nascent chain binding factors includ- ing NAC, SRP, and the Hsc70 co-chaperone St13. Engagement by St13 via its STI1 domain facilitates Hsc70 loading onto the mature domain early during biogenesis. The Hsc70-associated complex can recruit Hsp90 via the bridging co-chaperone Stip1. Hsp90 co-chaperones, which can also interact with the MTS, may exchange with St13 and re-model over time, with p23 favouring a stable substrate-Hsp90 complex due to p23 slowing the ATPse cycle of Hsp90 . Prolonged Hsp90 retention prevents mitochon- drial precursors from folding and prevents inappropriate interactions, thus keeping them in an import-competent state. All of the chaperone complexes during this process are likely to be import com- petent, with the Tom70 receptor (not shown) binding to Hsp70 or Hsp90. Transient exposure of the MTS, due its dynamic interaction with co-chaperones, would allow it to engage Tom20 or Tom22 (not shown), MTS receptors at the outer mitochondrial, to initiate import.

Analysis in the crosslinking assay (with the crosslinker in the MTS) of St13(ΔTPR) lacking the TPR domain showed that this mutant also lost substrate interaction (Fig. 6A). Affinity purification of St13(ΔTPR) showed that it purified as a single product, in contrast to St13 and St13(ΔSTI1), which co-purified with its known interaction partners Stip1, Hsc70, and Hsp90 (Fig. S5A). Of note, the crosslinking reaction with St13(ΔSTI1) showed increased crosslinking to Stip1 (Fig. S5B). This suggests that Stip1, a St13 homologue which has two STI1 domains and two TPR domains (Wang *et al*., 2022), can partially compensate for St13 function. These results suggest that St13 engages the MTS via its STI1 domain and that its interaction partners (Stip1, Hsc70 and Hsp90) are co-recruited and likely also interact with the nascent chain. A stable interaction with the substrate seems to rely on not only the STI1 domain, but also these interaction partners given that the interaction is lost upon deletion of the TPR domain that recruits them.Presumably, the avidity of a multi-valent interaction by different members of an otherwise dynamic complex stabilizes the substrate-(co)chaperone interactions. This would provide an explanation for why longer nascent polypeptides more readily crosslinked to St13 (Fig. 5F), as those would be better substrates for St13-Hsc70 complexes.

## Discussion

The early cytosolic events in mitochondrial protein biogenesis and their relationship to quality control remain mostly unresolved. In this study, we have systematically probed the molecular environment around the MTS, the most widely used signal for mitochondrial import, to reveal a set of co-chaperones for Hsc70 and Hsp90. In doing so, we defined a previously unappreciated role for the MTS in loading and retention of Hsc70 and Hsp90 onto mitochondrial precursors. This function becomes particularly important during import stress, when mitochondrial precursors are elevated and reside for prolonged periods in the cytosol. Hsp90 retains these precursors in an import-competent state, preventing their degradation and allowing import upon stress resolution. The ability to begin re-populating mitochondria with their residents immediately after stress may facilitate recovery.

Our findings lead to the following working model for how an MTS facilitates chaperone loading and retention on a mitochondrial precursor to maintain its import competence. As an MTS emerges from a translating ribosome, it seems to sample three nascent chain binding factors: (i) NAC, an exceptionally abundant triage factor that resides on most cytosolic ribosomes (Wiedmann *et al*., 1994; Gamerdinger *et al*., 2015; Gamerdinger and Deuerling, 2023); (ii) SRP, the ER targeting factor (Akopian *et al*., 2013); (iii) St13, a co-chaperone for Hsc70 (Li *et al*., 2013). The proximity to NAC is common to all nascent chains, and the SRP interaction is salt-sensitive, suggesting that it is sampling the nascent chain but not engaging it as a substrate. The St13 interaction occurs via its STI1 domain, whose shallow hydrophobic groove (Fry *et al*., 2021) would accommodate the hydrophobic face of an amphipathic MTS helix. This interaction is labile, consistent with the dynamic mode of interaction by other STI1 domains such as SGTA (Shao *et al*., 2017) and Ubiquilin family proteins (Itakura *et al*., 2016).

As more polypeptide emerges and is ultimately released from the ribosome, St13 recruits and facilitates loading of Hsc70. It does this by binding near the nucleotide-binding region of Hsc70, stabilizing its ADP-bound state, and inhibiting its ATPase cycle (Höhfeld *et al*., 1995; Li *et al*., 2013). Hsc70•ADP has high affinity for substrate (Rosenzweig *et al*., 2019), on which it lingers due to a slowed ATPase cycle. Prolonged retention of the St13-Hsc70 complex is presumably important for maintaining the precursor in an unfolded state until it is captured by the mitochondrial import receptors Tom22 and Tom20 at the outer mitochondrial membrane (Fig. 6). St13 is likely to be the long-sought "presequence binding factor" (PBF), a ∼50 kD protein of unknown identity that was observed in early work to interact with the presequence of pOTC (Murakami and Mori, 1990). Not only does St13 have an apparent migration of ∼50 kDa, but it cooperates with Hsc70 to engage pOTC more efficiently. This parallels the finding that addition of Hsc70 to pOTC-PBF complexes improved pOTC import into mitochondria in vitro (Murakami and Mori, 1990). Sti1, a homolog of St13 in yeast, interacts with yeast mitochondrial precursors and shows a synthetic growth phenotype with the mitochondrial import factors TOM20 and MIM1, further supporting a functional role in precursor import (Hoseini *et al*., 2016).

The substrate-St13-Hsc70 complex recruits Stip1, which recruits Hsp90 (Frydman and Höhfeld, 1997; Schopf *et al*., 2017; Rosenzweig *et al*., 2019; Wang *et al*., 2022). In the absence of St13, Stip1 may directly engage the MTS, presumably via its own STI1 domains. In this manner, the MTS-containing substrate becomes adorned with both the Hsc70 and Hsp90 systems. The dynamic interaction between the MTS and STI1 domains presumably facilitates exchange among Hsc70 and Hsp90 co-chaperones with affinity for the MTS. This would explain the observation that in total cytosol, the MTS can be photo-crosslinked to St13, Cdc37, p23, and to a lesser extent, Stip1. Although we do not have temporal information regarding the order of these interactions, studies of the Hsc70-Hsp90 system in folding cytosolic proteins suggest that substrates are progressively handed from the Hsc70 system to the Hsp90 system via the bridging factor Stip1 (Schopf *et al*., 2017; Rosenzweig *et al*., 2019; Wang *et al*., 2022).

The substrate-Hsp90 interaction is likely to be the final and relatively stable complex in this pathway. The stability of this interaction is probably facilitated by p23, which simultaneously interacts with the MTS and is thought to prolong Hsp90-substrate interactions (McLaughlin *et al*., 2006). Thus, both the Hsc70 and Hsp90 interactions are primarily via their ’holdase’ modes rather than the more dynamic folding modes. This prolongs the residence time of the chaperones on the mature domain and maintains the import competence of mitochondrial precursors. In both the Hsc70 and Hsp90 cases, interactions between the MTS and co-chaperones are relatively dynamic and context-dependent (i.e. can only occur when there is an additional interaction between the general chaperone and the mature part of the precursor). A dynamic interaction would periodically expose the MTS. Thus, once Tom70 at the outer mitochondrial membrane interacts with the general chaperone (either Hsc70 or Hsp90) (Young *et al*., 2003), the MTS would dynamically be accessible to import receptors such as Tom20 and Tom22 to initiate mitochondrial import (Brix *et al*., 1997, 1999; Su *et al*., 2022).

The substrate would be retained in an import-competent state throughout this sequence of events and hence capable of engaging the import machinery at any time. The ability to engage and use multiple MTS binding factors and two general chaperones explains why the system is highly robust to depletion or inhibition of one or more components, and why earlier genetic and biochemical strategies failed to identify a singular ’targeting factor’ analogous to SRP for the ER membrane. Only under conditions of prolonged cytosolic residence is a marked dependence on Hsp90 observed, the absence of which results in rapid degradation of many MTS-containing mitochondrial precursors (∼10% of all unimported precursors detected in the cytoplasm). This underscores the importance of Hsp90-mediated buffering of unimported mitochondrial proteins during stress.

Buffering by the Hsp90 system would not only prevent cellular toxicity by accumulated precursors, but also allows their rapid import without de novo synthesis when the stress has resolved. This may facilitate recovery from stress by allowing more rapid re-population of the mitochondrial proteome. A conceptually similar phenomenon of storing precursors for later import has been reported recently in yeast, where they are sequestered in reversible granules (Krämer *et al*., 2023). With prolonged stress, the precursors are degraded in mammalian cells, presumably as Hsp90 eventually cycles off these substrates or recruits a ubiquitin ligase such as CHIP (McDonough and Patterson, 2003). The pathway of degradation remains to be defined, but could also involve the Ubiquilin family of proteins (Itakura *et al*., 2016). The Ubiquilins not only contain STI1 domains that might engage the MTS (Fry *et al*., 2021), but also recruit ubiquitin ligase(s) to mark clients for degradation (Itakura *et al*., 2016). The SIFI complex could then recognize the pre-ubiquitinated precursor for polyubiquitination and targeting to the proteasome(Carrillo Roas *et al*., 2024; Grabarczyk *et al*., 2024; Yang *et al*., 2024).

### Materials and Methods Cell culture

The study used HEK293 Flp-In Trex-based cell lines. The parental cell line was obtained directly from the manufacturer (Life Technologies). All cells were cultured in Dulbecco’s Modified Eagle’s Medium (DMEM) with 10% Tetracycline-free Fetal Calf Serum (FCS). pOTC- GFP-P2A-RFP reporter was stably integrated into the FRT locus of HEK293 Flp-In Trex cells by co-transfection with Flp recombinase using the manufacturer’s protocol. Successfully modified cells were selected by culturing in media supplemented with 15 ug/ml blasticidin and 100 ug/ml hygromycin. For the induction of stably integrated transgenes, cells were treated with 1 ug/mL of doxycycline for the indicated time. CRISPR-Cas9 mediated knockout of ST13 in HEK293 Flp-In Trex cells was generated as described previously (Juszkiewicz and Hegde, 2017) using the following guide RNA: 5’- GCTATAGGAAATTTACCCTC-3’. Single cell-derived clones were screened for gene disruption by western blotting. All transient transfections of plasmids were performed with the TransIt 293 reagent (Mirus). siRNA silencing was typically for 3 days, using Silencer Select siRNAs (Life Technologies) transfected with Lipofectamine RNAiMAX (Life Technologies) according to the manufacturer’s protocol. Cell lines were routinely checked for mycoplasma contamination and verified to be negative.

### Antibodies and constructs

All antibodies, constructs, and gene blocks are listed in Table 1 below. The pOTC-GFP-P2A- RFP reporter construct was generated by insertion of the synthetically synthesized pOTC sequence (IDT) into the previously described pcDNA5-FRT/TO-GFP-P2A-RFP reporter backbone (Itakura *et al*., 2016) using a standard restriction enzyme-based cloning. Constructs for the transient expression of human ST13 were based on the pCMV vector. Synthetically synthesized ST13-1xFLAG (IDT) was inserted into the pCMV vector with Gibson Assembly (NEB). ΔTPR, ΔSTI, and all amber mutants were generated using inverse PCR and pCMV- St13-1xFLAG parental vector. In vitro translation reactions used synthetically synthesized gene blocks (IDT) encoding custom expression cassettes based on sp64 vector.

### Flow cytometry

Flow cytometry of the fluorescent reporter followed previously established protocols (Juszkiewicz and Hegde, 2017). Briefly, the stably integrated reporter was induced with 1 ug/ml doxycycline for the time indicated in the figure legends. When indicated, cells were co-treated with 100 nM valinomycin or 10 µM CCCP to inhibit mitochondrial import, 2 µM geldanamycin to inhibit Hsp90, 30 µM VER-155008 to inhibit Hsc70, and 10 µM MG132 to inhibit proteasome. After indicated time of induction (typically 6-8h), cells were washed with PBS, trypsinized, resuspended in PBS with 2% FCS, spun for 3 min at 5000 rpm in tabletop centrifuge at 4 °C and resuspended in ice-cold PBS. Cells were analyzed using LSRII instrument (Becton Dickinson) and data was processed using the FlowJo software. Each experiment used at least 20,000 GFP-positive events per each condition. The GFP:RFP ratio was plotted as a histogram. The histograms within any graph are directly comparable because the data were collected at the same time with the same detector settings, whilst the histograms from different graphs should not be compared as they could have been collected on different instruments and/or with different detector settings.

### Western blot analysis

For analysis of the total cellular protein, cells were washed with PBS and lysed with 100 mM Tris pH 8.0 with 1% SDS. Cell lysates were heated at 95 °C for 10 min and vortexed vigorously to shear the DNA. Samples were adjusted to the same concentration based on A280 values and 5xSDS sample buffer (500 mM Tris, 5% SDS, 50% glycerol, 500 mM DTT) was added to at least 1x final concentration. For the analysis of samples generated using in vitro translation, the same loading buffer was added directly to the sample of interest to at least 1x final concentration. Electrophoresis used 9% or 12% Tris-Tricine based gels. After electrophoresis, proteins were transferred to 0.2 um nitrocellulose membrane. Blocking and antibody incubations were for 1 h at room temperature or overnight at 4 °C with 5% non-fat powdered milk in PBS containing 0.1% Tween-20 (PBS-T). Detection employed HRP-conjugated secondary antibodies and SuperSignal West Pico Chemiluminescent Substrate (Thermo Fisher).

### *In vitro* transcription and translation in rabbit reticulocyte lysate

*In vitro* transcription was performed with SP6 polymerase using PCR products as the template as follows. The transcription reactions were conducted with 5-20 ng/ml template DNA in 40 mM HEPES pH 7.4, 6 mM MgCl2, 20 mM spermidine, 10 mM reduced glutathione, 0.5 mM ATP, 0.5 mM UTP, 0.5 mM CTP, 0.1 mM GTP, 0.5 mM m7G(5′)ppp(5′)G cap analog, 0.4-0.8 U/ml RNasin and 0.4 U/ml SP6 polymerase at 37 °C. In vitro translation in Rabbit Reticulocyte Lysate (RRL) was as described previously in detail (Sharma *et al*., 2010; Feng and Shao, 2018). In brief, translations were for 20-45 minutes at 32 °C. Translation reactions typically contained 33% by volume nuclease-treated RRL, 0.5 mCi/ml 35S-methionine, 20 mM HEPES, 10 mM KOH, 40 mg/ml creatine kinase, 20 mg/ml pig liver tRNA, 12 mM creatine phosphate, 1 mM ATP, 1 mM GTP, 50 mM KAc, 2 mM MgCl2, 1 mM reduced glutathione, 0.3 mM spermidine and 40 µM of each amino acid except methionine. The transcription reaction was added without further purification to 5% by volume to the translation reaction. Where indicated, 25 U/ml apyrase was added at the end of the reaction, followed by 15 min incubation at 32 °C to deplete ATP.

### Mitochondrial import assay

Mitochondrial import assay was performed using the *in vitro* translation system described above supplemented with freshly prepared semi-permeabilized (SP) cells with minor modifications of previously described methods (Itakura *et al*., 2016). In brief, HEK293 cells were plated on a 10 cm dish(es) ∼24h before the intended experiment. On the day of the experiment, ∼90% of confluent dishes containing cells were placed on ice, media were aspirated, and cells were gently washed with ice-cold PBS. To permeabilize cells, 2 ml of digitonin-based permeabilization buffer (20 mM HEPES pH 7.6, 100 mM KAc, 1.5 MgAc_2_, 0.005% purified digitonin) was added to each 10 cm dish. After 5 min incubation, the permeabilization buffer was aspirated, and cells were washed with ice-cold Physiological Salt Buffer (PSB: 20 mM HEPES pH 7.6, 100 mM KAc, 1.5 MgAc_2_). Cells were collected in PSB, spun again, and resuspended in 50 µl of PSB per each dish. 2 µl of SP cell suspension was used per each 10 µl (total) in vitro translation reaction prepared as described above. Additionally, the reaction mixtures were supplemented with 10 mM sodium succinate. Where indicated, 2 µM geldanamycin was added to inhibit Hsp90, and 1 µM valinomycin was added to inhibit mitochondrial import. All the components were added on ice, and in vitro translation and mitochondrial import reactions were performed in 32 °C water bath for 1h.

### Immunoprecipitation and affinity purification

To immunoprecipitate (IP) in vitro translated mitochondrial precursors (or crosslinked adducts) and associated proteins under native conditions, IVT reactions were first diluted 5-fold with 1xRNC buffer (50 mM HEPES pH 7.6, 100 mM KAc, 1.5 MgAc_2_) followed by addition of indicated antibodies and packed resins (either Protein A or G for samples containing antibodies or Strep-Tactin for direct purification of precursors via TST). All the antibodies used in this study are listed in Table 1. IPs were incubated at 4 °C for 3h with end-over-end rolling. Resins were then washed 5 times with 1xRNC buffer. Samples incubated with antibodies were eluted directly with SDS sample buffer and analysed by SDS-PAGE and autoradiography. Samples incubated with Strep-Tactin resin were first eluted with 1xRNC buffer containing 50 mM biotin for 30 min on ice, then mixed with SDS sample buffer and analysed by SDS-PAGE and autoradiography. IP under denaturing conditions used similarly prepared IVT and photo- crosslinked samples which were first denatured by addition of 100 mM Tris pH 8.0 and 1% SDS, followed by 5 min of heating at 95 °C. Samples were then diluted 10-fold to reduce SDS concentration using IP buffer (50 mM HEPES pH 7.6, 100 mM NaCl, 10 mM MgAc_2_, 0.5% Triton X-100). Antibodies and resins were added as described above and samples were incubated for 3h with end-over-end rolling at 4 °C. Samples were washed thrice with IP buffer prior to elution with SDS buffer (antibody-containing samples) or IP buffer containing 50 mM biotin for 30 min on ice (Strep-Tactin containing samples). All samples were analyzed by SDS-PAGE and autoradiography.

### Purification of St13 mutants

ΔST13 HEK293 Flp-In Trex cells were transfected with indicated ST13 constructs as described above (used 10 µg of plasmid per 10 cm dish). After 48 h of expression, cells were washed with ice-cold PBS, and collected by gentle pipetting in ice-cold PBS. After a 3 min spin at 4 °C at 5000 rpm in a tabletop centrifuge, cell pellets were resuspended in hypotonic buffer (10 mM HEPES pH 7.6, 10 mM KAc, 1.5 mM MgAc_2_, 2 mM DTT). Cells were swelled on ice for 15 min and then lysed mechanically by passing through 26G needle attached to a 1 ml syringe 30 times. Buffer concentrations were then adjusted either to physiological salt (20 mM HEPES pH 7.6, 100 mM KAc, 2 mM MgAc_2_), or high salt (40 mM HEPES pH 7.6, 400 mM KAc, 2 mM MgAc_2_), and lysates were spun at 4 °C for 15 min at max speed in tabletop centrifuge to remove the debris. Supernatants were mixed with FLAG-M2 resin and incubated for 90 min at 4 °C with end-over-end rolling. The beads were washed with either physiological salt or high salt buffer four times. FLAG-tagged proteins were eluted with 0.25 mg/ml FLAG peptide in the physiological salt buffer for 30 min at room temperature. Purified proteins were analyzed by SDS-PAGE or used directly in crosslinking experiments as indicated.

### Preparation of cytosolic fractions from HEK cells

WT HEK293 Trex cells were treated with siRNA targeting ST13, STIP1, or non-targeting control for 3 days, whilst ST13 KO (ΔST13) cells were transiently transfected with plasmids encoding ST13 mutants as described above, and expression of transgenes was allowed for 48h. After washing with PBS, cells were collected in ice-cold PBS by gentle pipetting. Cells were then spun at 4 °C for 3 min at 5000 rpm in a tabletop centrifuge. Cell pellets were resuspended in hypotonic buffer (10 mM HEPES pH 7.6, 10 mM KAc, 1.5 mM MgAc_2_, 2 mM DTT) and incubated on ice for 15 min. Mechanical lysis was performed by passing through 26G needle attached to 1 ml syringe 30 times. Buffer conditions were then adjusted to 20 mM HEPES pH 7.6, 100 mM KAc, 2 mM MgAc_2_. Cell lysates were spun at 4 °C for 15 min at max speed in a tabletop centrifuge to remove the debris. Lysate concentration was measured on nanodrop using A280 readings and adjusted to 10 mg/ml with the same buffer. Fresh cell lysates were used directly in downstream experiments as indicated in legends.

### Site specific photo-crosslinking with p-Benzoyl-L-phenylalanine (BpA)

BpA, a UV-activated crosslinker, was incorporated into defined sites of an in vitro translation product by amber suppression as previously described (Chfiven *et al*., 2003; Lin *et al*., 2020). In short, the standard RRL translation system was supplemented with 5 μM B. stearothermophilus tRNATyr, 0.25 μM E. coli derived tyrosyl-tRNA synthetase mutated to accept BpA (Chin *et al*., 2003), and 0.1 mM BpA. Following any subsequent biochemical manipulations as described in the figure legends, the samples were irradiated on ice ∼10 cm away from a UVP B-100 series lamp (UVP LLC) for 10 minutes. The samples were either analyzed directly or subjected to subsequent fractionation or immunoprecipitation as described in the legends.

### Site-specific photo-crosslinking with 3’-azibutyl-N-carbamoyl-lysine (AbK)

Site-specific incorporation of AbK crosslinker into the STI1 domain of St13 followed previously established protocol (O’Donnell *et al*., 2020). Briefly, ΔST13 cells were co-transfected with plasmids encoding ST13 modified with amber codons at positions indicated in Figure 6C, and a pAS-Pyl-AF plasmid encoding the *Methanosarcina mazei* pyrrolysyl-tRNA synthetase carrying Y306A and Y384F mutations and tRNAPylCUA pair. To achieve comparable expression of amber mutants, each 10 cm dish of ΔST13 cells was transfected with 4 µg of pAS-Pyl-AF plasmid and 100 ng of no-amber control ST13, 500 ng of F335amber-ST13, 1000 ng of M345amber-ST13, 1000 ng of Q336amber-ST13, or 200 ng of N356amber-ST13 plasmids. Expression of ST13 mutants was for 2 days in the presence of 0.5 mM AbK. Lysates with AbK-incorporated ST13 were then generated as described above.

### Sucrose gradient fractionations

20 µl IVT reaction of pOTC-TST with BpA incorporated at position F3 was layered atop 200 µl of 5-25% sucrose gradients prepared at least 1h before by layering 40 µl fractions of 25%, 20%, 15%, 10% and 5% sucrose solutions in 1xRNC by hand. Sucrose gradients were spun for 2h 20 min at 55 000 rpm in 4 °C using TLS-55 rotor and Optima Max-XP centrifuge (Beckman Coulter) set up to slowest acceleration and deceleration speeds. 11 fractions (20µl each) were then collected starting from the top of the gradient, and each individual fraction was crosslinked with UV as described above. Crosslinked samples were mixed with SDS sample buffer and analysed SDS-PAGE and autoradiography.

### Purification of ribosome-nascent chain complexes

Ribosome-nascent chain complexes (RNCs) of 102-mer pOTC with or without BpA incorporated into positions indicated in figure legends were generated in an RRL-based in vitro translation system (described above) using truncated mRNA as a template (Sharma *et al*., 2010; Feng and Shao, 2018). Briefly, 50 µl in vitro translation reactions were incubated at 32°C for 10 min, followed by transfer on ice to stop translation. Samples were then adjusted to high salt (500 mM KAc and 7.5 mM MgAc_2_) or diluted with an equivalent amount of 1xPSB, and layered atop a 150 µl of 20% sucrose cushion prepared in 50 mM pH 7.6 HEPES-based buffer supplemented with high salt (500 mM KAc, 7.5 mM MgAc_2_) or physiological salt (100 mM KAc, 5 mM MgAc_2_). Samples were spun for 1h at 100 000 rpm at 4 °C using TLA120.1 rotor and Optima Max-XP centrifuge (Beckman Coulter). The supernatant was aspirated, and RNC pellets were resuspended in ∼25 µl of 1xPSB, RRL, HEK lysates (prepared as described above), or purified proteins (purified as described above). When indicated, nascent chains were released from the ribosome by an additional 5 min incubation at 32 °C in the presence of 1 mM puromycin. The samples were then UV-crosslinked as described above and either analyzed directly or subjected to subsequent immunoprecipitation as indicated in the legends. All nascent chain-containing samples were supplemented with 0.01M EDTA and RNAse A (0.1 mg/ml) and incubated for 15 min on ice to remove the tRNA immediately before SDS-PAGE.

### Identification of pOTC interaction partners

Two replicates of in vitro translation reactions (1 ml each) of pOTC-TST or GFP-TST were affinity purified via TST using StrepTactin resin as described above. One replicate of the washed resin was eluted with SDS-PAGE sample buffer for electrophoresis and visualization of recovered proteins with SYPRO Ruby stain (Fig. 1A). The other replicate of the washed resin was subjected to on-bead trypsin digestion and analysis by mass spectrometry as follows. Proteins on the resin were reduced with 2 mM DTT in 2 M urea, 50 mM Tris pH8 and sequencing grade trypsin (Promega) was added to a final concentration of 5 ng/µl. After 1 h incubation at 25°C, supernatants were then transferred to clean tubes. Beads were washed twice with 2 M urea in 50 mM Tris, and the washes were combined with the corresponding supernatants. Next, supernatants were alkylated with 4 mM iodoacetamide, and an additional 0.1 µg of trypsin was added to the samples before incubation overnight at 25°C. Samples were acidified with formic acid (FA) and desalted using home-made C18 (3M Empore) stage tip packed with Poros oligo R3 resin (Thermo Scientific). Bound peptides were eluted with 30- 80% MeCN/0.5% FA. Eluates were partially dried in a Speed Vac (Savant) for LC-MS-MS analysis as described below.

### Quantitative mass spectrometry of cytosolic extracts

The experiment used eighteen 10 cm dishes (6 per condition) of HEK 293 Trex cells in total. Cells treated as indicated in the legend to Fig. 2 were extracted with 500 µl of digitonin-based permeabilization buffer (20 mM HEPES pH 7.6, 100 mM KAc, 1.5 MgAc2, 0.01% purified digitonin) per each plate. The cells were separated from the extracted cytosol by centrifugation in a tabletop microcentrifuge (10 min at maximum speed) at 4°C. Cytosolic extracts from two 10 cm dishes were pooled together for each condition to form one technical replicate. In total, three technical replicates for each of the three conditions were used for downstream analysis. The cytosolic extracts in 0.01% digitonin were reduced with 5 mM dithiothreitol (DTT) at 37°C for 1 h and alkylated with 10 mM iodoacetamide in the dark at room temperature for 30 min. Excess iodoacetamide was quenched with 5 mM DTT for 10 minutes. Samples were first digested with Lys-C (Promega) for 4 h at 25°C followed by trypsin (Promega) incubated overnight, at 30°C. Digestion was stopped by the addition of formic acid (FA) to a final concentration of 0.5% and centrifuged at 16,000 x g for 5 min to remove any particulate matter.

Supernatants were desalted using home-made C18 stage tips (3M Empore), filled with 2 mg of Oligo R3 (Thermo Scientific) resin. Stage tips were equilibrated with 80% acetonitrile (MeCN)/0.5 %FA followed by 0.5%FA. Bound peptides were eluted with 30-80% MeCN/0.5% FA and lyophilized.

The lyophilized peptides from each condition were resuspended in 200 mM Hepes, pH 8.5. TMT 10-plex reagent (Thermo Scientific) reconstituted in anhydrous MeCN according to manufacturer’s instruction was added at a ratio of 1:3 (peptides:TMT reagent) so that peptides from each condition were labelled with a distinct TMT tag for 60 min at room temperature. The labelling reaction was stopped by incubation with 5% hydroxylamine for 30 min. Labeled peptides were combined into a pooled sample and partially dried to remove MeCN in a SpeedVac (Thermo Scientific). After this, the sample was desalted as before and the eluted peptides were lyophilized.

TMT labelled peptides were subjected to off-line High-Performance Liquid Chromatography (HPLC) fractionation, using XBridge BEH130 C18, 5 µm, 2.1 x 150 mm (Waters) column with XBridge BEH C18 5 µm Van Guard Cartridge, connected to an Ultimate 3000 Nano/Capillary LC System (Dionex). Peptides were separated with a gradient of 1-90% B (A: 5% MeCN/10 mM ammonium bicarbonate, pH 8; B: MeCN/10 mM ammonium bicarbonate, pH 8, [9:1]) in 1 h at a flow rate of 250 µl/min. Eluted peptides was collected at 1 min/ fraction for 60 min, then combined into 15 fractions and lyophilized. Dried peptides were resuspended in 1% MeCN/0.5% FA, desalted using C18 stage tips and partially dried down in a Speed Vac for LC-MS-MS analysis as described below.

### Liquid Chromatography-Mass spectrometry Analysis

All samples were analysed by LC-MS/MS using a fully automated Ultimate 3000 RSLC nano System (Thermo Fisher Scientific) fitted with a 100 μm x 2 cm PepMap100 C18 nano trap column and a 75 μm × 25 cm, nanoEase M/Z HSS C18 T3 column (Waters). Peptides were separated using a binary gradient consisting of buffer A (2% MeCN, 0.1% formic acid) and buffer B (80% MeCN, 0.1% formic acid), at a flow rate of 300 nL/min. Eluted peptides were introduced directly via a nanoFlex ion source into a Q Exactive Plus (for protein identification in Fig. 1A) or an Orbitrap Eclipse (for the TMT labelled samples for Fig. 2) mass spectrometer (Thermo Fisher Scientific).

Data dependent acquisitions were carried out on the protein identification samples (Fig. 1A). MS1, full-scan (m/z 380-1600) with a resolution of 70K, followed by MS2 acquisitions of the 15 most intense ions with a resolution of 17.5K and NCE (normalized collision energy) of 27%. MS1 target values of 1e6 and MS2 target values of 5e4 were used. The isolation window was set at 1.5 m/z and dynamic exclusion for 40s.

Orbitrap Eclipse data acquisition of TMT labelled fractions (Fig. 2) were performed in real-time database search (RTS) with synchronous-precursor selection (SPS) -MS3 analysis. MS1 spectra were acquired using the following settings: Resolution=120K; mass range=400- 1400m/z; AGC target=4e5; MaxIT=50ms and dynamic exclusion was set at 60s. MS2 analysis were carried out with HCD activation, ion trap detection, AGC=1e4; MaxIT=50ms; CE (fixed) =35% and isolation window =0.7m/z. RTS of MS2 spectrum was set up to search uniport human proteome, with fixed modifications cysteine carbamidomethylation and TMT 10-plex at N-terminal and Lys residue. Met-oxidation was set as variable modification. Missed cleavage=1 and maximum variable modifications=2. In MS3 scans, the selected precursors were fragmented by HCD and analyze using the orbitrap with these settings: Isolation window=1 m/z; CE (fixed) =55, orbitrap resolution=50K; scan range=110-500 m/z; MaxIT=200ms and AGC=1e5.

### Analysis of mass spectrometry data

Raw data from the affinity purification samples for protein identification (Fig. 1A) were searched against Uniprot Fasta protein database (Mammalia was selected, downloaded 2016), using the Mascot search engine (Matrix Science, v2.4). Database search parameters were set with a precursor tolerance of 10 ppm and a fragment ion mass tolerance of 0.1 Da. A maximum of two trypsin missed cleavages was allowed. Carbamidomethyl cysteine was set as static modification while oxidized methionine and acetylation of protein N-terminal were specified as variable modifications. Scaffold (version 4, Proteome Software Inc.) was used to validate MS/MS-based peptide and proteins identifications. Protein identifications were accepted if they were greater than 90% probability assigned by the Protein Prophet algorithm and contained at least 2 identified peptides, and are shown in Table S1.

Raw data from the TMT experiment (Fig. 2; Table S2) was processed using MaxQuant (Cox and Mann, 2008) with the integrated Andromeda search engine (v.1.6.17.0). MS/MS spectra were quantified with reporter ion MS3 from TMT10plex experiments and searched against Homo sapiens Fasta protein database (UP000005640_9606_1gene=1prot, downloaded on August 2022). Carbamidomethylation of cysteines was set as fixed modification, while methionine oxidation and protein N-terminal acetylation were set as dynamic modifications. Tryptic digestion up to 2 missed cleavages were allowed. MaxQuant output file, proteinGroups.txt was then analyzed with Perseus software (v 1.6.15.0). After uploading the matrix, the data was filtered to remove identifications from reverse database, identifications with modified peptide only, and common contaminants. The resulting matrix was log 2 transformed, normalized by median of columns and only kept those rows with all 9 valid values. Then this matrix was exported as a text file for further data analysis as displayed in Fig. 2 and Fig. S1, with data shown in Table S2.

## Acknowledgments

We are grateful to the members of the Hegde lab for helpful discussions. This work was supported by UK Medical Research Council grant MC_UP_A022_1007 (R.S.H.).

---

**Figure S1.**
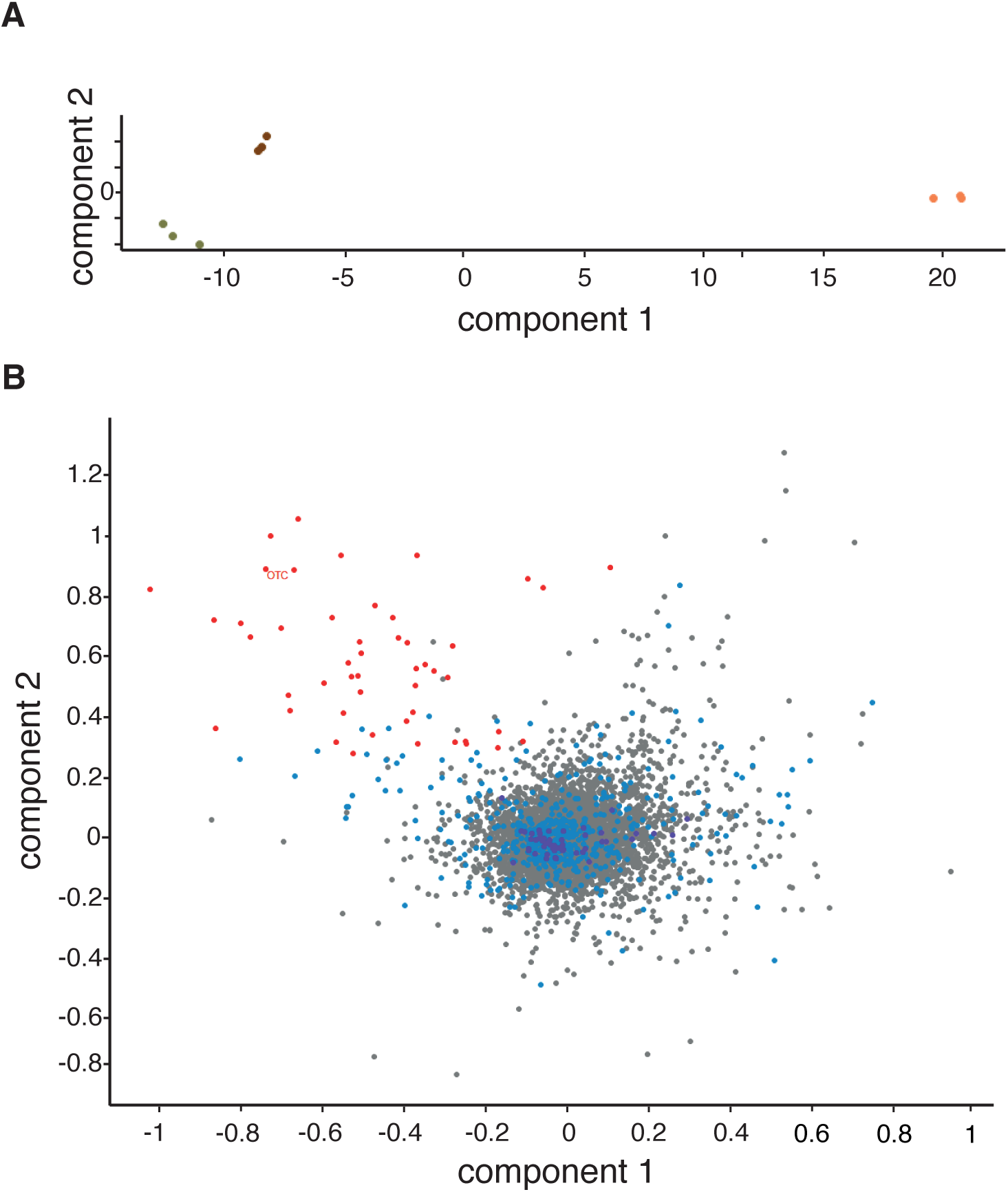
Hsp90 broadly buffers mitochondrial precursor degradation during import stress. (A) Principal component analysis (PCA) of the three technical replicates for each of the three biological conditions (depicted in three different colors) analyzed in the mass spectrometry experiment presented in Figure 2. Note clustering of the replicates indicative of high reproducibility of the results. **(B)** Principal component analysis (PCA) of the dataset from the the TMT experiment depicted in Figure 2C. Note that Hsp90 dependent proteins (red dots) tend to cluster together in the upper left quadrant of the plot.

**Figure S2.**
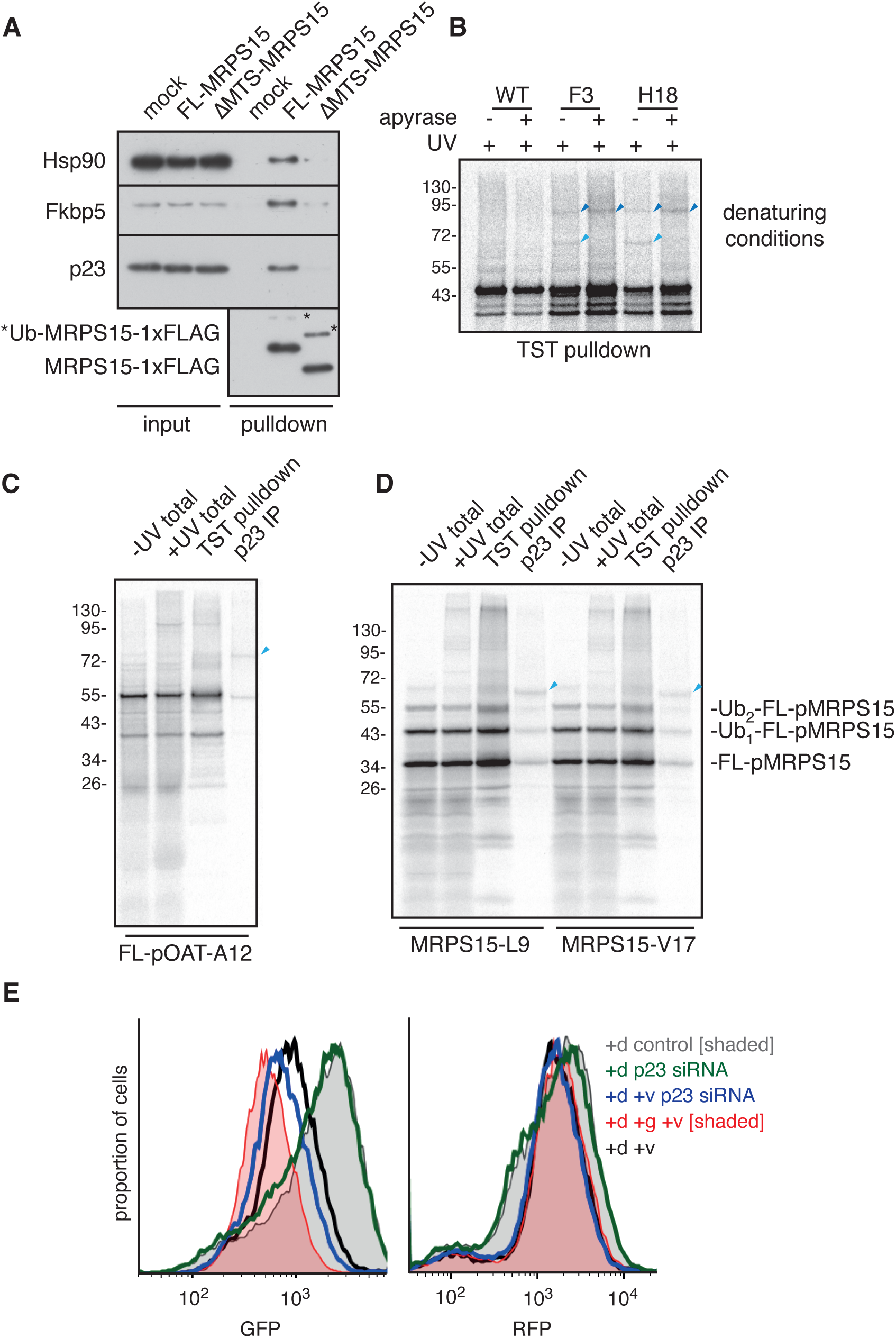
MTS facilitates Hsp90 retention. (A) FL or ΔMTS MRPS15 was translated in RRL in the presence or absence of 2 μM geldanamycin and affinity purified under native conditions via a C-terminal FLAG tag. Samples eluted with FLAG peptide were analyzed by immunoblotting alongside input samples. Ubiquitinated MRPS15 is marked with asterisk. **(B)** FL-pOTC-TST with BpA crosslinker at positions F3 or H18 was produced in RRL. Samples were then UV irradiated either prior to or after ATP depletion with apyrase. All samples were affinity purified via the TST tag on the substrate and analyzed by SDS-PAGE and autoradography. Note the disappearence of the smaller crosslink (light blue arrow) upon ATP depletion. **(C)** FL-pOAT-TST with BpA crosslinker in the MTS (position A12) was produced in RRL and UV irradiated. Samples were then fully denatured and affinity purified using Streptactin (TST pulldown) or anti-p23 antibody. **(D)** MRPS15-TST with BpA at positions L9 or V17 was produced in RRL and UV irradiated. Samples were then fully denatured and affinity purified using Streptactin (TST pulldown) or anti-p23 antibody. **(E)** Histograms representing GFP or RFP signal from the experiment presented in Figure 3F. Note the selective decrease in GFP signal (without an appreciable change in RFP signal) in cells treated with siRNA against p23 (blue traces).

**Figure S3.**
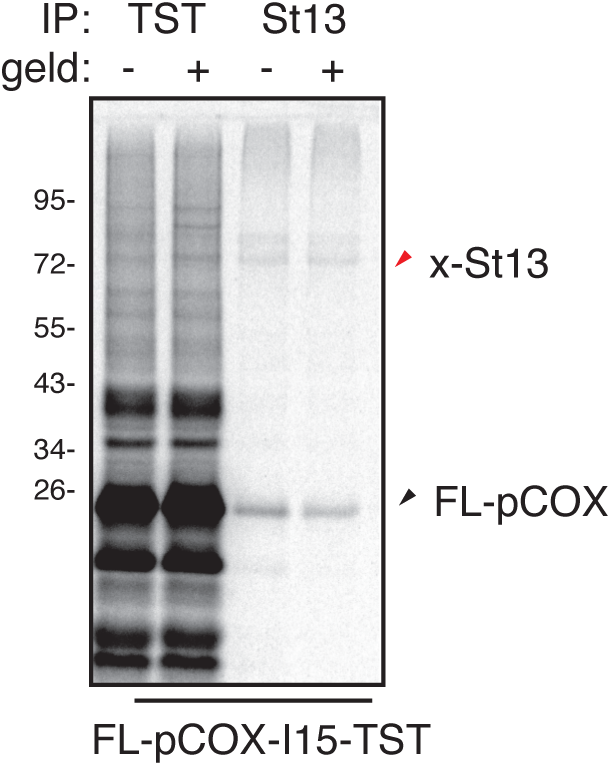
St13 interact with the MTS of pCOXIVI1. BpA-containing pCOXIVl1 was produced in RRL in the presence or absence of geldanamycin and crosslinked with UV. Samples were then fully denatured and affinity purified using Streptactin (TST pulldown) or anti-St13 antibody. pCOX-St13 crosslink is marked with red arrow.

**Figure S4.**
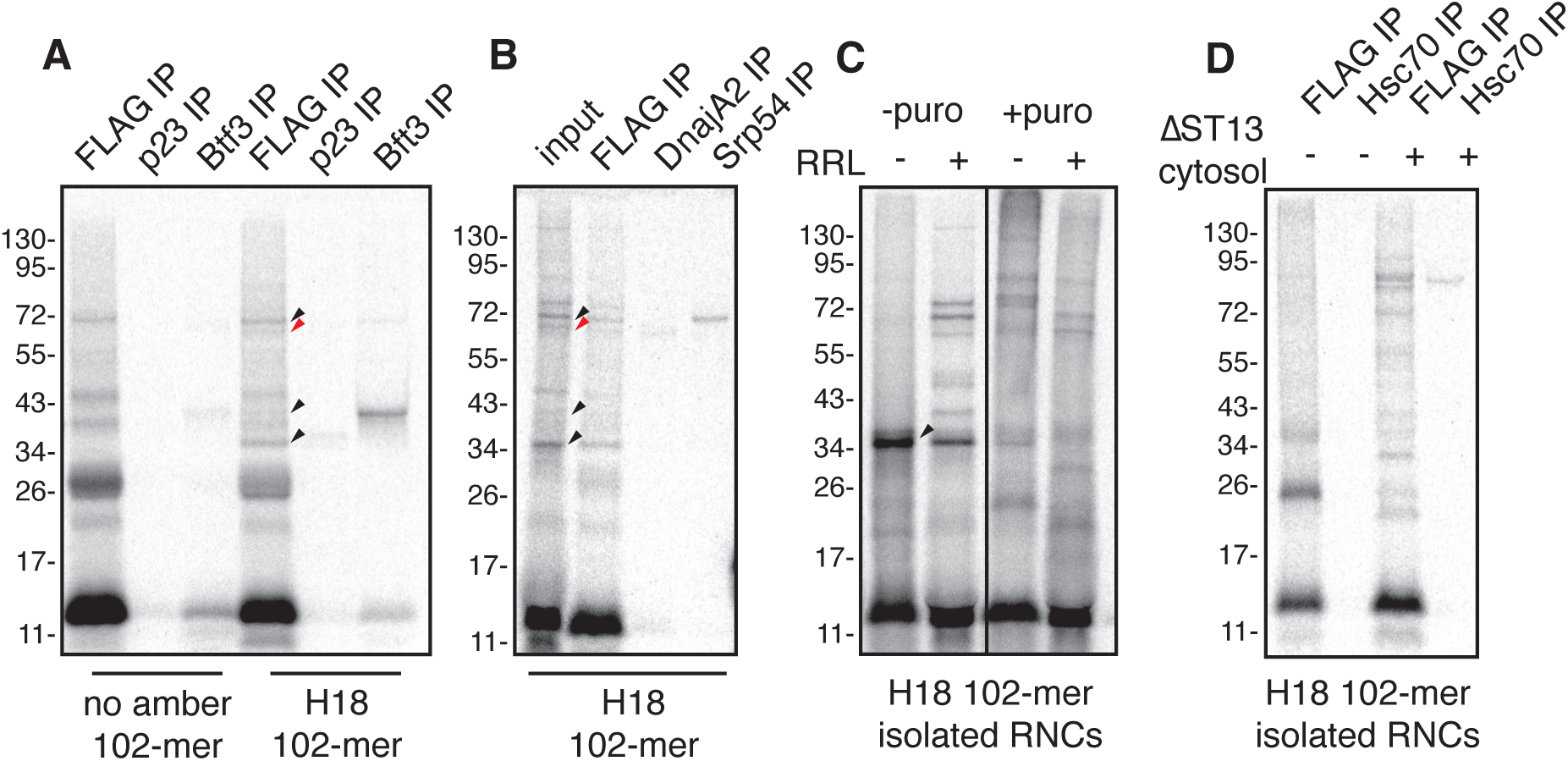
St13 engages the MTS co-translationally and is retained after ribosome release. (A) 102-mer pOTC ribosome-nascent chain complexes (RNCs; see Figure 5A) with or without BpA at position H18 were produced in RRL. After UV crosslinking, samples were denatured and immunoprecipitated (IP) with antibodies against FLAG (substrate), p23 or Btf3. **(B)** An experiment as in panel A, but with IP using antibodies against DnajaA2 and Srp54. **(C)** 102-mer pOTC RNCs with BpA at positon H18 were synthesized in RRL and separated from bulk cytosol under high-salt conditions via centrifugation. The resuspended RNCs were mixed with RRL or buffer where indicated, released from ribosomes using puromycin where indicated, and subjected to UV irradiation. Note that of the several crosslinks observed in panel A, only the smallest of them is retained after RNCs are isolated in high salt (arrowhead). The lost crosslinks are restored if the RNCs are mixed with RRL. **(D)** 102-mer pOTC RNCs with BpA at positon H18 were synthesized in RRL and separated from bulk cytosol under high-salt conditions via centrifugation. The resuspended RNCs were mixed with or without cytosolic extract from ΔST13 HEK cells and subjected to UV irradiation. Samples were then subjected to IP with antibodies against the substrate (FLAG) or Hsc70.

**Figure S5.**
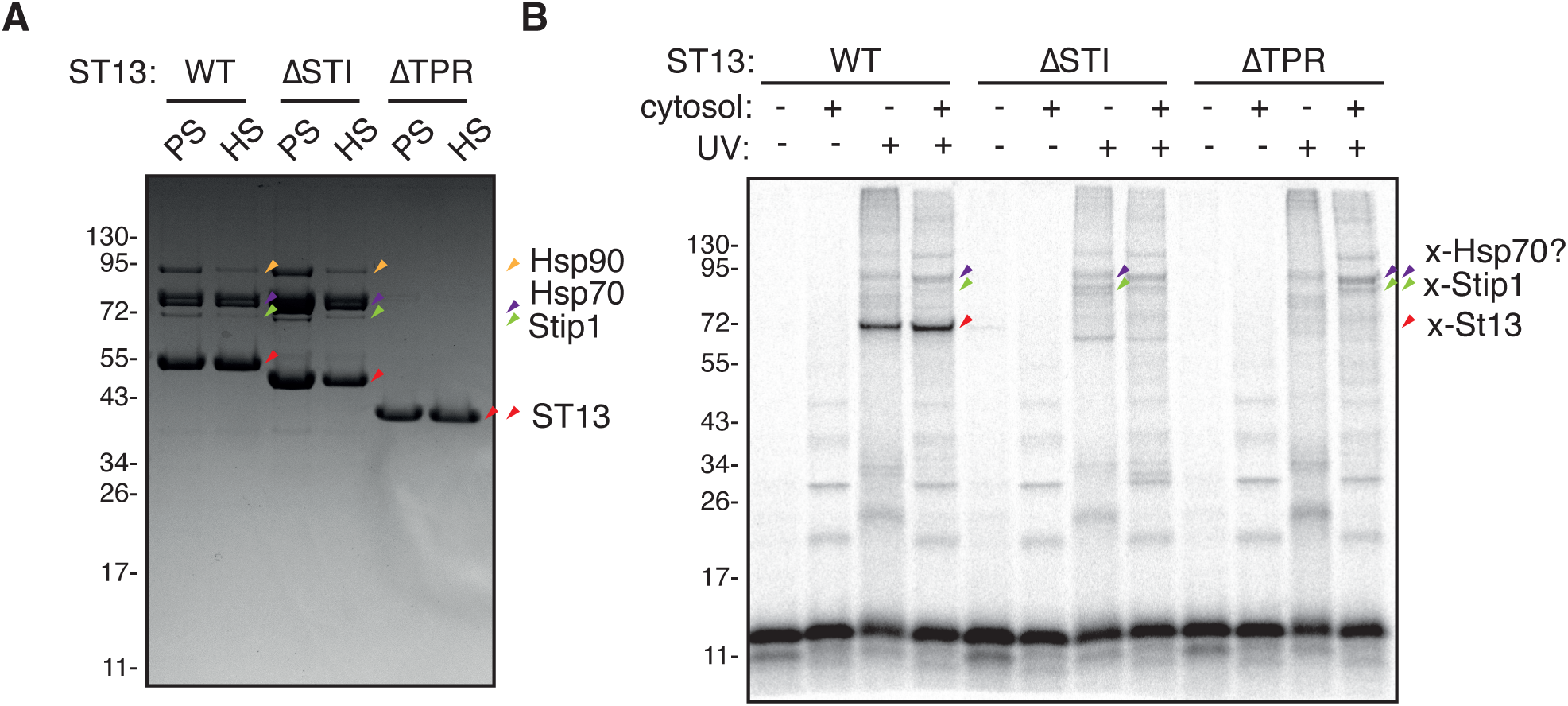
St13 engages MTS via its STI1 domain. (A) FLAG-tagged ST13 recombinant proteins (WT, ΔSTI1, ΔTPR) were produced in HEK cells and purified under physiological salt (PS) or high salt (HS) conditions. Purified proteins were analyzed by SDS-PAGE and Coomassie staining. ST13 mutants are marked with red arrows, whereas known interacting partners are labeled as follows: Hsc70 is marked with purple arrows, Stip1 with green arrows and Hsp90 with orange arrows. Note the disapperance of interacting partners in the ΔTPR mutant. **(B)** Ribosome-nascent chain complexes (RNCs) of 102-mer pOTC were produced in RRL and purified by centrifugation through a sucrose cushion in high salt buffer. Stripped RNCs were then resuspended in buffer containing the indicated purified proteins from the high salt condition in (A) alone or together with cytosolic extracts from ΔST13 cells. Note the strongest signal for St13 crosslink when WT protein was added (even in the absence of the cytosol).

